# Regulation of replication origin licensing by ORC phosphorylation reveals a two-step mechanism for Mcm2-7 ring closing

**DOI:** 10.1101/2023.01.02.522488

**Authors:** Audra Amasino, Shalini Gupta, Larry J. Friedman, Jeff Gelles, Stephen P Bell

**Affiliations:** Howard Hughes Medical Institute, Department of Biology, Massachusetts Institute of Technology, Cambridge, MA 02139, USA; Department of Biochemistry, Brandeis University, Waltham, MA 02454, USA

**Author notes:** Co-corresponding authors: Stephen P. Bell, Jeff Gelles.

## Abstract

Eukaryotic DNA replication must occur exactly once per cell cycle to maintain cell ploidy. This outcome is ensured by temporally separating replicative helicase loading (G1 phase) and activation (S phase). In budding yeast, helicase loading is prevented outside of G1 by cyclin-dependent kinase (CDK) phosphorylation of three helicase-loading proteins: Cdc6, the Mcm2-7 helicase, and the origin recognition complex (ORC). CDK inhibition of Cdc6 and Mcm2-7 are well understood. Here we use single-molecule assays for multiple events during origin licensing to determine how CDK phosphorylation of ORC suppresses helicase loading. We find that phosphorylated ORC recruits a first Mcm2-7 to origins but prevents second Mcm2-7 recruitment. Phosphorylation of the Orc6, but not of the Orc2 subunit, increases the fraction of first Mcm2-7 recruitment events that are unsuccessful due to the rapid and simultaneous release of the helicase and its associated Cdt1 helicase-loading protein. Real-time monitoring of first Mcm2-7 ring closing reveals that either Orc2 or Orc6 phosphorylation prevents Mcm2-7 from stably encircling origin DNA. Consequently, we assessed formation of the MO complex, an intermediate that requires the closed-ring form of Mcm2-7. We found that ORC phosphorylation fully inhibits MO-complex formation and provide evidence that this event is required for stable closing of the first Mcm2-7. Our studies show that multiple steps of helicase loading are impacted by ORC phosphorylation and reveal that closing of the first Mcm2-7 ring is a two-step process started by Cdt1 release and completed by MO-complex formation.

**Significance Statement:** Each time a eukaryotic cell divides (by mitosis) it must duplicate its chromosomal DNA exactly once to ensure that one full copy is passed to each resulting cell. Both under-replication or over-replication result in genome instability and disease or cell death. A key mechanism to prevent over-replication is the temporal separation of loading of the replicative DNA helicase at origins of replication and activation of these same helicases during the cell division cycle. Here we define the mechanism by which phosphorylation of the primary DNA binding protein involved in these events inhibits helicase loading. Our studies identify multiple steps of inhibition and provide new insights into the mechanism of helicase loading in the uninhibited condition.

## Introduction

Eukaryotic cells have evolved mechanisms to ensure that exactly one complete copy of their genome is created during each mitotic cell cycle (reviewed in (Arias and Walter, 2007)). There is a strong selective advantage to this precise control: either under-replicating or over-replicating the genome has deleterious consequences including cell death, genome instability, and cancer and developmental abnormalities in animals (reviewed in (Gaillard et al., 2015)). To ensure complete replication, each eukaryotic chromosome initiates replication from many origins of replication, which are present in excess across each chromosome (Prioleau and MacAlpine, 2016). This abundance of origins facilitates complete DNA replication, but also presents a challenge to the cell: ensuring that none of the many origins initiate replication more than once during a cell cycle.

Eukaryotic cells prevent origins from repeated initiation during a cell cycle by temporally separating two key replication events: helicase loading and helicase activation (Siddiqui et al., 2013). Helicase loading (also known as origin licensing or pre-RC formation) is restricted to the G1 phase of the cell cycle and defines all potential origins of replication (Bell and Labib, 2016). Activation of the loaded helicases only occurs during S phase (Tanaka et al., 2007; Zegerman and Diffley, 2007). Importantly, once the cell enters S phase and throughout G2 and M phases, helicase loading is actively inhibited.

Four proteins work together to load helicases at origins of replication: the origin recognition complex (ORC), Cdc6, Cdt1, and the core engine of the replicative DNA helicase, the Mcm2-7 complex (reviewed in (Bell and Labib, 2016; Stillman, 2022). ORC recognizes origin DNA and recruits Cdc6 to complete a protein ring encircling the DNA (Fig. 1A, step 1, Feng et al., 2021; Schmidt et al., 2022). The ORC-Cdc6-DNA complex recruits and binds to the C-terminal face of the ring-shaped Mcm2-7 complex bound to Cdt1 to form the ORC-Cdc6-Cdt1-Mcm2-7 (OCCM) intermediate (Fig. 1A step 2, Randell et al., 2006; Yuan et al., 2017). The Mcm2-7 ring is recruited to the DNA in an open state, with a gap between the Mcm2 and Mcm5 subunits (the Mcm2-5 gate, (Bochman and Schwacha, 2008; Samel et al., 2014) that allows DNA access to its central channel. Shortly after OCCM formation, Cdc6 and then Cdt1 depart in an ordered manner (Fig. 1A, steps 3-4; Ticau et al., 2015). Closing of the first Mcm2-7 ring occurs concomitant with Cdt1 departure and requires Mcm2-7 ATP hydrolysis (Ticau et al., 2017). After the first Mcm2-7 is loaded, ORC can “flip” relative to the initial DNA binding site and first-Mcm2-7/DNA complex (Gupta et al., 2021) and a new interaction is established between ORC and Mcm2-7 in which ORC binds to the opposite, N-terminal face of the first Mcm2-7 complex and to an inverted DNA binding site (Fig. 1A, step 5; the “MO complex”; Miller et al., 2019). Cdc6 and a second Mcm2-7/Cdt1 are recruited to the MO complex in the opposite orientation to the first (Fig. 1A, steps 6-7). The second Mcm2-7 rapidly forms interactions with the first Mcm2-7 via their N-terminal domains and MO interactions are lost (Fig. 1A step 8, Gupta et al., 2021; Miller et al., 2019; Ticau et al., 2015). Sequential release of Cdc6 followed by Cdt1 and ORC results in closing of the second Mcm2-7 and the formation of a tightly interacting head-to-head dimer of two Mcm2-7 complexes that encircle double-stranded DNA (Fig. 1A, steps 9-10; Evrin et al., 2009; Remus et al., 2009; Ticau et al., 2017).

**Figure 1:**
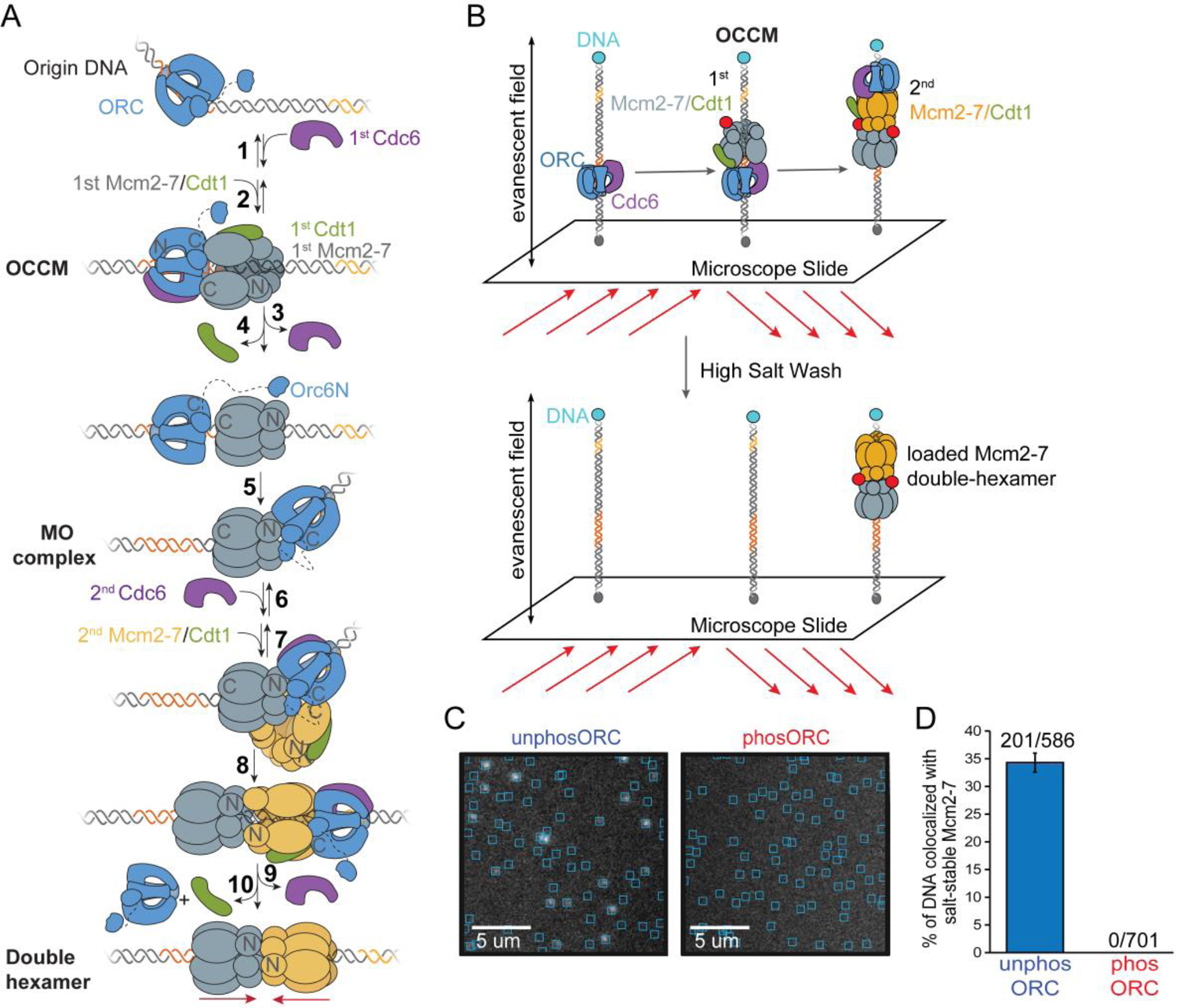
ORC phosphorylation inhibits the formation of salt-stable Mcm2-7 complexes on DNA in single-molecule experiments. **A.** Model of the events of the helicase loading. See text for description. First Mcm2-7 complexes are shown as gray and second Mcm2-7 complexes are shown as yellow. Dark orange region on DNA represents the initial high-affinity ORC binding site (ACS), the yellow region of DNA represents a 2^nd^ inverted, weaker ORC binding site (B2). **B.** Schematic of experiment. Fluorescent-dye-labeled (red) Cdt1-Mcm2-7^4SNAPJF646^, ORC, and Cdc6 were incubated in a chamber with surface-tethered, fluorescent dye-labeled (blue) DNA^AF488^ containing *ARS1* (red DNA). 20-minute reactions were followed by a high-salt wash to remove intermediates. Protein and DNA fluorescence at the slide surface was monitored using TIRF microscopy. **C.** Representative micrograph segments showing Mcm2-7^4SNAPJF646^ fluorescence colocalized with sites of individual DNA molecules (cyan boxes) after a high-salt wash for reactions including either unphosORC (left) or phosORC (right). **D.** Percent (± SEM) of DNA molecules associated with fluorescent Mcm2-7 after a high-salt wash.

In the budding yeast *S. cerevisiae*, helicase loading occurs at specific DNA sequences and is inhibited outside of G1 by cyclin-dependent kinase (CDK) activity. Budding yeast origins of replication include one strong (ACS) and at least one oppositely-oriented and weaker DNA-binding sites (B2) for ORC (Coster and Diffley, 2017; Marahrens and Stillman, 1992; Wilmes and Bell, 2002). CDK independently inhibits helicase loading by phosphorylating three of the four helicase-loading proteins: Mcm2-7, Cdc6, and ORC. CDK phosphorylation of Mcm2-7 induces the nuclear export of Mcm2-7 (and any associated Cdt1) molecules that have not been loaded onto origin DNA during G1 (Labib et al., 1999; Liku et al., 2005; Nguyen et al., 2000). Cdc6 phosphorylation leads to its ubiquitin-modification and degradation (Drury et al., 2000, 1997; Elsasser et al., 1999). In contrast to Mcm2-7 and Cdc6, phosphorylated ORC remains in the nucleus bound to origins of replication throughout the cell cycle (Aparicio et al., 1997). These findings suggest that ORC phosphorylation intrinsically inhibits one or more event(s) during helicase loading. This conclusion is supported by *in vitro* helicase-loading assays that show that CDK phosphorylation of ORC, but not Cdc6 or Mcm2-7, inhibits this event (Chen and Bell, 2011; Frigola et al., 2013; Phizicky et al., 2018).

Ensemble biochemical assays narrow down which steps of helicase loading could be impacted by ORC phosphorylation: phosphorylated ORC is capable of binding origin DNA and recruiting all the other helicase-loading proteins but fails to form the Mcm2-7 double hexamer (Frigola et al., 2013). Single-molecule assays for helicase-loading enable individual observation of many additional steps relative to ensemble assays. These include initial recruitment of the first and second Mcm2-7/Cdt1 complexes, sequential release of Cdc6 and Cdt1 after each Mcm2-7 recruitment, MO complex formation, and Mcm2-7 ring closure around DNA (Gupta et al., 2021; Ticau et al., 2017, 2015).

To understand how ORC phosphorylation inhibits helicase loading, we performed single-molecule helicase-loading assays containing either phosphorylated or unphosphorylated ORC. These studies demonstrate that multiple steps of helicase loading are impacted by ORC phosphorylation, the most important of which is the prevention of MO complex formation, an event that we demonstrate is required to form a stably-bound closed-ring state of the first Mcm2-7 complex.

## Results

### Single-molecule experiments verify that ORC phosphorylation inhibits helicase loading

We used established single-molecule (SM) assays to observe intermediates in helicase loading in the presence of phosphorylated ORC (phosORC) or unphosphorylated ORC (unphosORC) (Ticau et al., 2015). Phosphorylated ORC was prepared by incubating Clb5-Cdk1 (hearfter, CDK) and ATP with ORC. Phosphorylation was terminated by adding the B-type-CDK-inhibitor Sic1 (Schwob et al., 1994), preventing CDK action during subsequent helicase-loading events. We labeled different pairs of helicase-loading proteins to evaluate distinct steps in the helicase-loading pathway. Importantly, all the fluorescently-labeled proteins used have been shown to be functional in previous ensemble and single-molecule (SM) helicase-loading assays (Gupta et al., 2021; Ticau et al., 2017, 2015). AlexaFluor-488 end-labeled DNA containing the *ARS1* origin of replication was used for Colocalization Single Molecule Spectroscopy (CoSMoS; Friedman et al., 2006; Hoskins et al., 2011) to evaluate interactions between origin DNA and the helicase-loading proteins.

To confirm that phosphorylated ORC is unable to load helicases in the SM setting, we used Mcm2-7 fluorescently labeled at the N-terminus of Mcm4 (Mcm2-7^4SNAPJF646^) to visualize Mcm2-7 DNA-binding events. To select for loaded Mcm2-7 molecules, at the end of each reaction we performed a high-salt wash that releases helicase-loading intermediates from DNA but retains loaded Mcm2-7 proteins (Fig. 1B, Donovan et al., 1997; Randell et al., 2006; Ticau et al., 2015). Consistent with previous ensemble assays (Chen and Bell, 2011; Frigola et al., 2013; Phizicky et al., 2018), SM helicase-loading assays with phosORC showed no high-salt-resistant Mcm2-7-DNA association (Fig. 1C and 1D).

### ORC phosphorylation slows OCCM formation

We next asked whether ORC phosphorylation alters the rate of initial recruitment of the Mcm2-7/Cdt1 complex on the pathway to form the OCCM. Previous ensemble assays showed both unphosORC and phosORC facilitated OCCM formation in the presence of ATPγS (Frigola et al. 2013). However, since ATPγS traps helicase loading at the OCCM step, these end-point studies may not have detected changes to the rate of OCCM formation. Using the SM helicase-loading assay, we asked if phosORC supported OCCM formation in the presence of ATP and, if so, whether the kinetics of this event were altered.

To measure the rate of OCCM formation, we determined the time from the start of the reaction to the first Mcm2-7 association with each DNA molecule (Fig. 2A, Friedman and Gelles, 2015). Because the first Mcm2-7-Cdt1 complex is recruited to DNA only after ORC and Cdc6 have bound, stable first Mcm2-7 recruitment is equivalent to OCCM formation (Fig 1A, step 2; Aparicio et al., 1997; Remus et al., 2009; Seki and Diffley, 2000; Ticau et al., 2015). ORC phosphorylation resulted in a 59% reduction in the rate of first Mcm2-7-Cdt1 recruitment (*k*_a_= 0.0047 ± 0.0003 s^−1^ with unphosORC vs. *k*_a_ = 0.0020 ± 0.0002 s^−1^ with phosORC; Figure 2B, Supp. Table 1). Thus, in the presence of ATP, ORC phosphorylation does not completely prevent OCCM formation but does significantly reduce its rate. This change may contribute to, but cannot by itself explain, the complete loss of helicase loading observed in Fig. 1.

**Figure 2:**
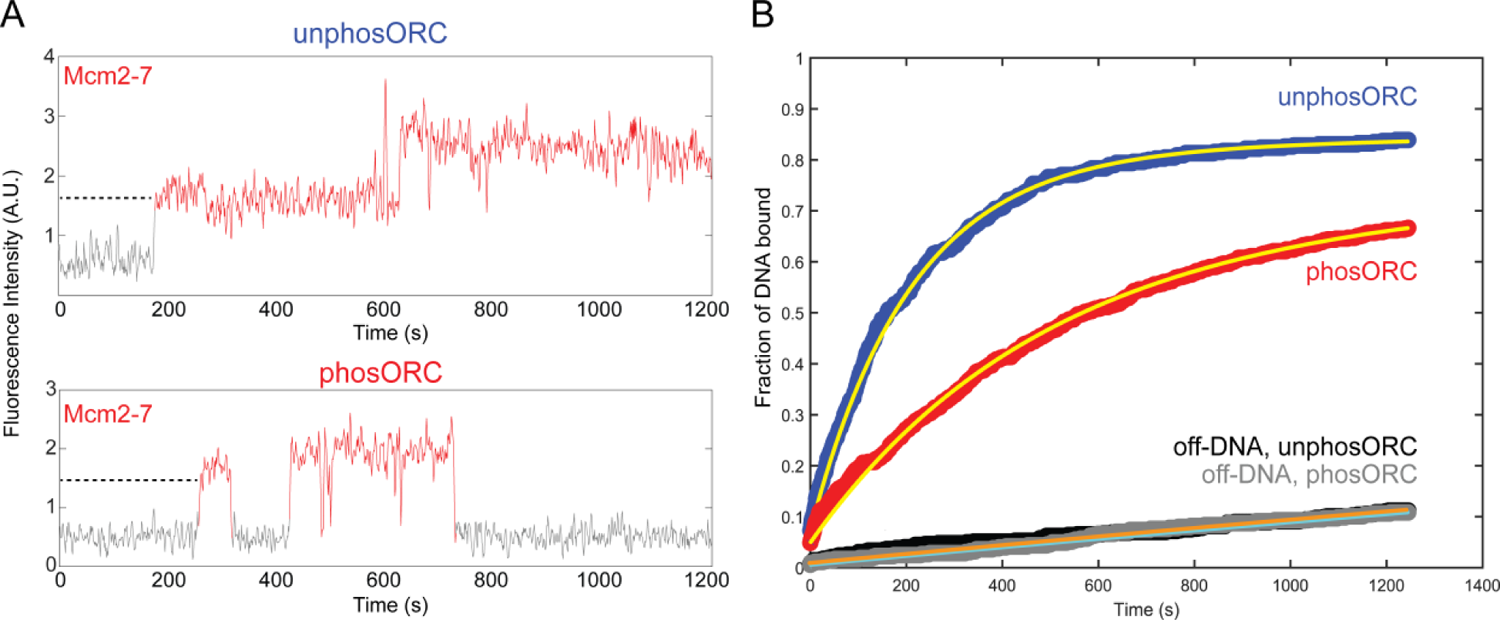
Phosphorylated ORC reduces the rate of OCCM formation. **A.** Example fluorescence records showing Mcm2-7^4SNAPJF646^ associating with single DNA molecules in experiments using either unphosORC (top) or phosORC (bottom), taken from the experiments in Fig. 1. Red color denotes times during which Mcm2-7^4SNAPJF646^ colocalizes with the DNA; grey denotes times with no Mcm2-7^4SNAPJF646^ colocalization. Dotted lines indicate the measured time interval before the first Mcm2-7 bound to each DNA molecule. **B.** Determining rates of Mcm2-7 association. Cumulative fraction of DNA molecules that have associated with at least one Mcm2-7 in the presence of unphosORC (blue) or phosORC (red) is plotted as a function of time after the start of the experiment. Data from control non-DNA locations on the slide surface from the same experiments (unphosORC, black; phosORC, gray) are also shown. Yellow lines show fits of the data to an exponential model (see Methods) yielding the fit parameters shown in Supp. Table 1. Orange (unphosORC) and Cyan (phosORC) lines show the exponential fit of background binding to non-DNA sites.

### ORC phosphorylation inhibits recruitment of the second Mcm2-7

We next examined the impact of ORC phosphorylation on recruitment of second Mcm2-7 complexes. As in previous studies (Gupta et al., 2021; Ticau et al., 2017, 2015), with unphosORC we frequently observe sequential long-lived associations of two Mcm2-7 complexes (Fig. 3A, corresponding to Fig. 1A, steps 2 and 7). In contrast, when ORC was phosphorylated, although we observed frequent long-lived first Mcm2-7 association with DNA, we did not observe stable second Mcm2-7 binding (Fig. 3B). Quantitatively, ORC phosphorylation reduced the fraction of DNAs with long-lived (> 15 s) 1^st^ Mcm2-7 associations approximately two-fold, most likely due to reduced OCCM formation (Fig. 3B). More importantly, phosphorylation entirely abolished long-lived 2^nd^ Mcm2-7 associations (Fig. 3C). Although we did observe occasional short-lived (< 15 s) 2^nd^ Mcm2-7 associations when ORC was phosphorylated, even these were rare (5/204 1^st^ Mcm2-7 events). Together, these findings show that ORC phosphorylation inhibits double-hexamer formation at a step between recruitment of the first and second Mcm2-7 complexes.

**Figure 3:**
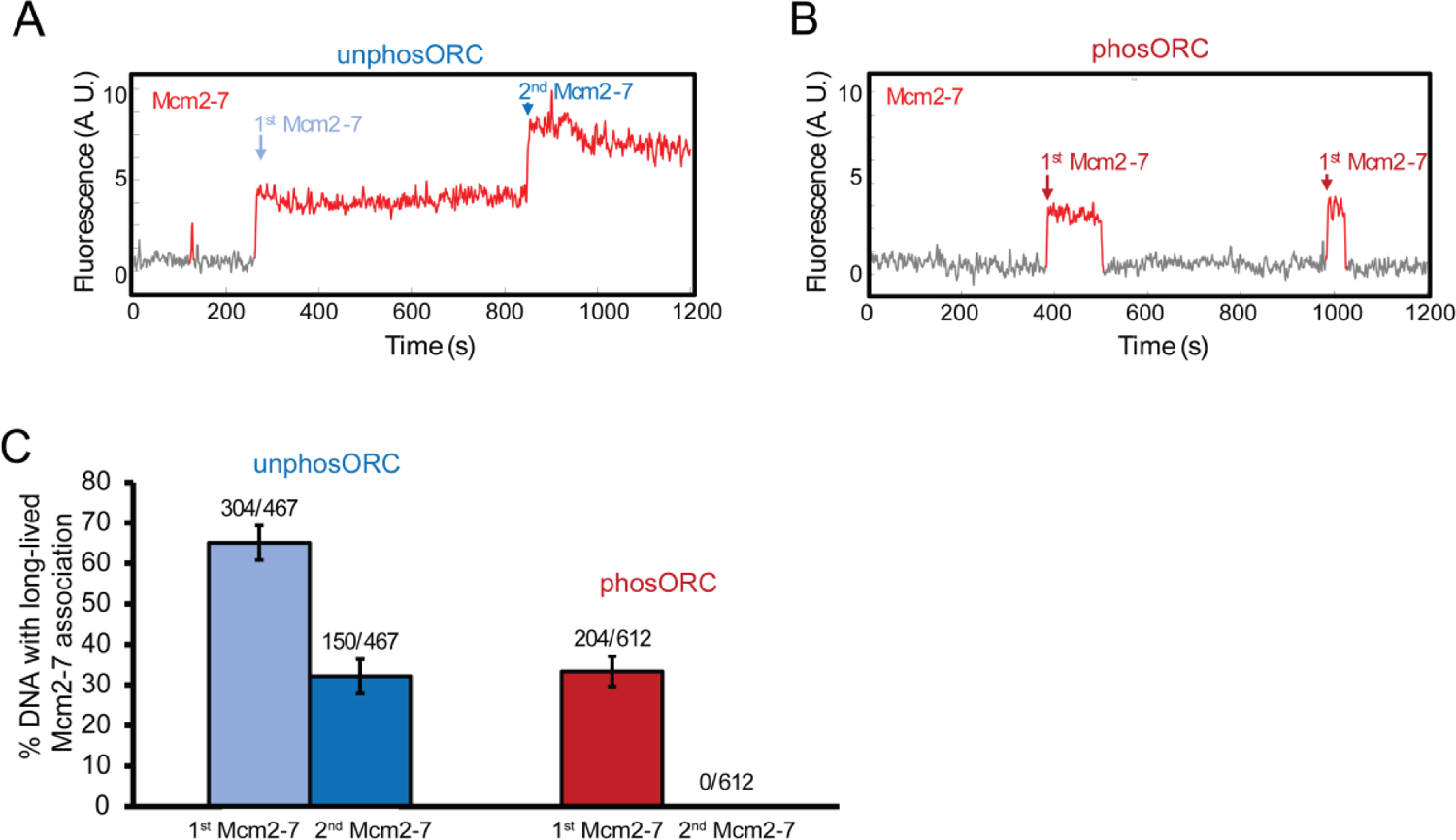
ORC phosphorylation reduces 1^st^ Mcm2-7-DNA associations and abolishes stable recruitment of 2^nd^ Mcm2-7 complexes. **A.** Representative record of Mcm2-7^4SNAPJF646^ associations with an individual DNA molecule in the presence of unphosORC (experiment same as in Fig. 1), plotted as in Fig. 2A. Additional example records are shown in Supp. Fig. 1A. **B.** Representative record of Mcm2-7^4SNAPJF646^ associations with an individual DNA molecule in the presence of phosORC. Two single (1^st^) Mcm2-7 association events were observed. Additional example records are shown in Supp. Fig. 1B. **C.** Percentage (± SEM) of DNA molecules exhibiting long-lived (i.e., >15s) 1^st^ Mcm2-7 and 2^nd^ Mcm2-7 binding to individual DNA molecules. A 2^nd^ Mcm2-7 binding event is defined as a second Mcm2-7 binding while the first Mcm2-7 is still present. Only such overlapping long-lived associations have the potential to form loaded Mcm2-7 double-hexamers (Ticau et al., 2017, 2015). DNA molecules that had multiple events, (i.e., as in B) were counted once using the longest event.

Together, our initial findings identify a key segment of the helicase-loading pathway that is fully inhibited by ORC phosphorylation. Although ORC phosphorylation results in delays in OCCM formation, it does not prevent it (Fig. 2), consistent with previous ensemble ATPγS experiments showing that OCCM formation still occurs when ORC is phosphorylated (Chen and Bell, 2011; Frigola et al., 2013; Phizicky et al., 2018). The absence of long-lived 2^nd^ Mcm2-7 associations in the presence of phosORC (Fig. 3) indicates that phosphorylation inhibits helicase loading prior to or at the step of second Mcm2-7 recruitment. To further elucidate the events altered by ORC phosphorylation, we focused our subsequent experiments on the events after recruitment of the 1^st^ Mcm2-7 and up to the arrival of the 2^nd^ Mcm2-7 (Fig. 1A, steps 3-7).

### ORC phosphorylation inhibits second Cdc6 recruitment

The next step in helicase loading after OCCM formation is release of Cdc6 (Fig. 1A, step 3 (Ticau et al., 2015). To address whether Cdc6 release is altered by ORC phosphorylation, we performed the SM helicase-loading assay using fluorescently-labeled Cdc6^SORT549^ and Mcm2-7^4SNAPDY649^-Cdt1 and unlabeled unphosORC or phosORC (Fig. 4A). We used arrival of the first Mcm2-7 with DNA to mark the time of OCCM formation and measured how long Cdc6 was retained after this event (Fig. 4B, Supp. Fig. 2A, B). Both unphosORC and phosORC yielded similar distributions of Cdc6 dwell times after first Mcm2-7 arrival (Fig. 4C). However, in the presence of phosORC we never observed recruitment of a second Cdc6 while the first Mcm2-7 remained on DNA (0/64). In contrast, a second Cdc6 is consistently observed before recruitment of a second Mcm2-7 when ORC is unphosphorylated (38/42, Fig4B, the four 2^nd^ Mcm2-7 events that are not preceded by a 2^nd^ Cdc6 are consistent with incomplete fluorescent labeling of Cdc6). We conclude that ORC phosphorylation does not alter the rate of Cdc6 dissociation from the OCCM but prevents a later step involved in or required for recruitment of a second Cdc6.

**Figure 4:**
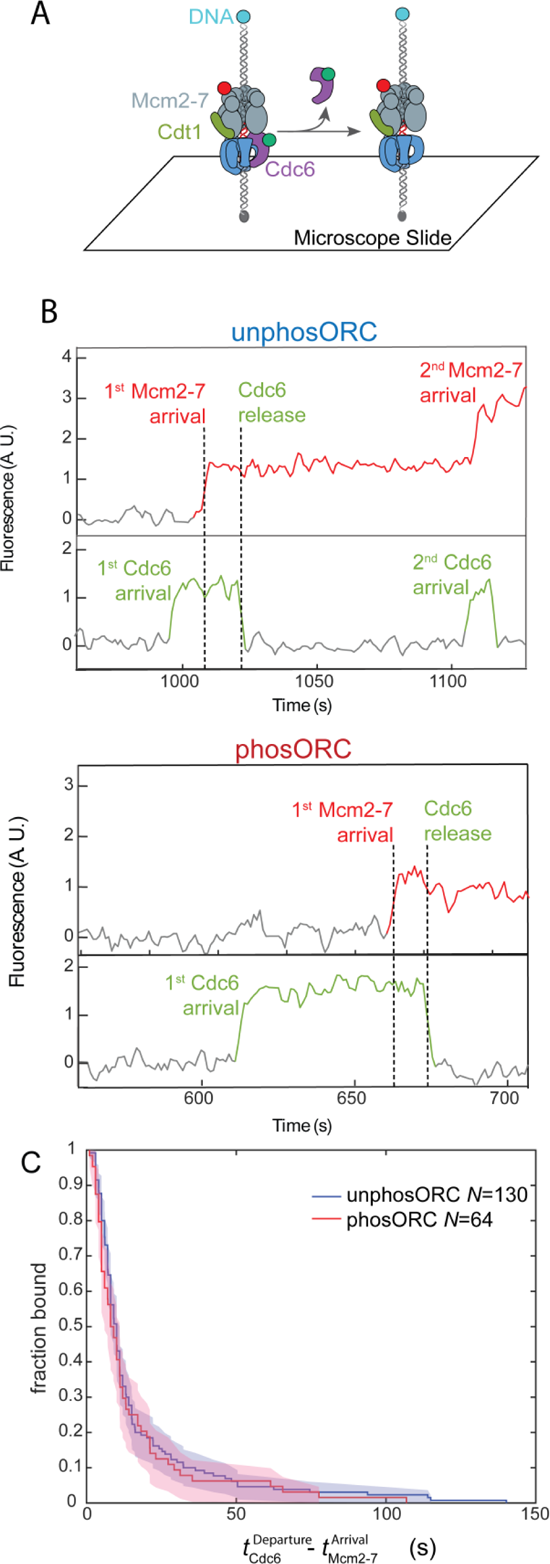
Cdc6 departure from the OCCM is not affected by ORC phosphorylation. **A.** Schematic of Cdc6 release experiment, illustrated as in Fig. 1B. **B.** Representative traces of labeled Mcm2-7^4SNAPDY649^ (red) and Cdc6^SORT549^ (green) binding to one DNA molecule in the presence of unlabeled unphosORC (top) or phosORC (bottom) and plotted as in Fig. 2A. The vertical dashed lines represent the beginning (Mcm2-7 arrival) and end (Cdc6 release) of the dwell interval of Cdc6 following the time of first Mcm2-7 association as plotted in Fig. 4C (this interval represents the duration of the OCCM). Additional examples are shown in Supp. Fig. 2. **C.** The fraction of DNAs that retain Cdc6 following first Mcm2-7 association (i.e., after OCCM formation) vs time in reactions with unphosORC (blue) and phosORC (red) plotted as cumulative survival functions. Shaded areas represent the 95% confidence intervals for each curve.

### ORC phosphorylation increases non-productive Cdt1 dissociation

During successful helicase loading, Cdc6 release is followed by release of Cdt1 and Mcm2-7 ring closure (Fig. 1A, steps 3 and 4). Events in which Mcm2-7 is retained on the DNA after Cdt1 release (“Cdt1-only release” events, Fig. 5A, right; Ticau et al., 2017, 2015) are on pathway to productive double-hexamer formation. In addition to these potentially productive events, previous studies observed an alternative, non-productive pathway after initial Cdt1-Mcm2-7 association: simultaneous release of the recruited Cdt1-Mcm2-7 complex (“simultaneous Cdt1-Mcm2-7 release”, Fig. 5A, left, Ticau et al., 2015). When Cdt1-Mcm2-7 dissociate simultaneously, Mcm2-7 has a characteristically shorter dwell time than when Cdt1 dissociates independently from Mcm2-7 (Ticau et al., 2015).

**Figure 5:**
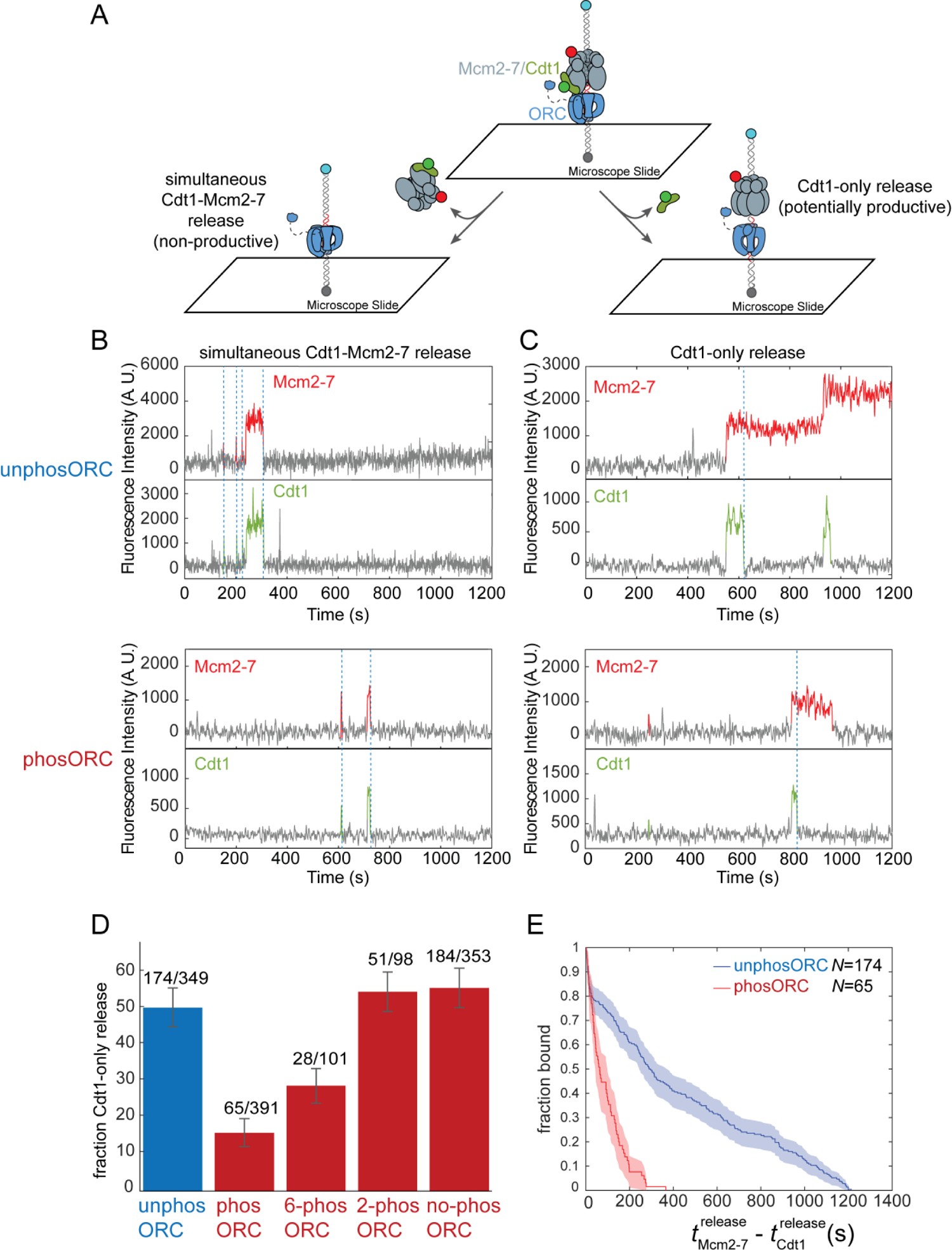
ORC phosphorylation increases the fraction of non-productive Mcm2-7-Cdt1 recruitment events. **A.** Schematic of experiment to monitor the two Cdt1 dissociation pathways. Left: Cdt1 and the associated Mcm2-7 release from the DNA simultaneously (non-productive). Right: Cdt1 releases from the DNA prior to Mcm2-7 (potentially productive). **B.** Example fluorescence records showing non-productive Mcm2-7^4SORTJF646^ (red) and Cdt1^SORTDY549^ (green) simultaneous dissociation (dashed lines) from a single DNA (left pathway in A). Experiments used unphosORC (top) or phosORC (bottom). Colored intervals indicate presence of fluorescent protein colocalized with DNA. Additional examples are shown in Supp. Fig. 3A and B. **C.** Example fluorescence records showing potentially productive Mcm2-7^4SORTJF646^ (red) and Cdt1^SORT549^ (green) in which Cdt1 dissociates from DNA before the corresponding first Mcm2-7 (dashed lines; right pathway in A). Experiments used unphosORC (top) or phosORC (bottom). Additional examples are shown in Supp. Fig. 3C and D). **D.** The percentage (± SEM) of first Cdt1 dissociation events that followed the productive pathway (Cdt1 release without Mcm2-7 release) is shown for wild-type ORC without CDK treatment (blue), or CDK-treated (red) wild-type ORC or mutant ORC constructs. Mutants allow phosphorylation of Orc6 alone (6-phosORC), Orc2 alone (2-phosORC), or neither subunit (no-phosORC). The impact of each mutant on phosphorylation was confirmed (Supp. Fig. 3G). Only first Mcm2-7 data is included in this analysis. **E.** The fraction of 1^st^ Mcm2-7 molecules that remain on DNA (as defined in Fig. 3A and 3B) after Cdt1 dissociation in experiments with unphosORC (blue) and phosORC (red) are plotted as cumulative survival functions. Shaded areas represent 95% confidence intervals.

To determine whether ORC phosphorylation impacts the choice of Cdt1-release pathways, we performed the SM helicase loading assay with fluorescently-labeled Cdt1^SORT549^ and Mcm2-7^4SNAPJF646^ (Fig. 5A). Regardless of ORC phosphorylation, we observed both simultaneous (Fig. 5B, Supp. Fig. 3A and 3B) and Cdt1-only (Fig. 5C, Supp. Fig. 3C and 3D) release events. However, ORC phosphorylation changed the fraction of OCCM complexes that were resolved by the Cdt1-only release pathway. In the absence of ORC phosphorylation 47 ± 5% OCCM complexes proceeded through the Cdt1-only release pathway, but when ORC was phosphorylated only 15 ± 4% of OCCMs followed this pathway (Fig. 5D; first and second bars). Although ORC phosphorylation decreases the percentage of Cdt1-only release events, the kinetics of the two release pathways are not significantly changed (Supp. Fig. 3E and 3F). Together, these data indicate that, in addition to reducing the rate of OCCM formation (Fig. 2), phosORC inhibits helicase loading by increasing the proportion of OCCM complexes that undergo simultaneous Mcm2-7-Cdt1 release and are, therefore, non-productive.

Because two ORC subunits, Orc2 and Orc6, are phosphorylated by CDK and both contribute to the prevention of helicase loading (Chen and Bell, 2011; Nguyen et al., 2001), we asked whether phosphorylation of one or both subunits decreased use of the potentially productive Cdt1-only release pathway. To this end, we prepared ORC constructs incorporating mutations that eliminate CDK phosphorylation of the mutated subunits (Supp. Fig. 3G, Nguyen et al 2001) and used the mutant constructs in the Cdt1-release assay (Fig. 5D). Phosphorylation of Orc6 alone (6-phosORC) showed use of the Cdt1-only release pathway (27 ± 4%) that was between unphosphorylated and fully phosphorylated ORC (47 ± 5% and 15 ± 4%, respectively). In contrast, phosphorylation of only Orc2 (2-phosORC) was indistinguishable from unphosORC (Fig. 5D). Importantly, when both Orc2 and Orc6 CDK phosphorylation sites were eliminated (no-phosORC), CDK treatment caused no change in pathway choice, indicating that there are no other CDK phosphorylation sites that independently affect Cdt1-release pathway selection (Fig. 5D; compare no-phosORC with unphosORC). When not treated with CDK, all three of the ORC phospho-site mutants showed rates of productive pathway use similar to WT unphosphorylated ORC (Supp. Fig. 3H). Thus, these mutations affect ORC function only by preventing phosphorylation.

In addition to its effect on pathway choice, ORC phosphorylation also strongly reduces the dwell times of Mcm2-7 molecules retained after Cdt1 release. Reactions with unphosORC resulted in Mcm2-7 complexes remaining on the DNA for long durations after Cdt1 release (Fig. 5E; median (*t*_Mcm2−7_^release^ − *t*_Cdt1_^release^) = 584 s). This median value may underestimate the actual dwell times of these Mcm2-7 complexes because 49% ± 7% (94/191) of the measured dwell times were artificially censored by the end of the experiment. In contrast, when ORC is phosphorylated, the Mcm2-7 complexes remaining on DNA after Cdt1 release have much shorter dwell times (Fig. 5E; median (*t*_Mcm2−7_^release^ − *t*_Cdt1_^release^) = 92 s) with only rare (5/209) censored events.

### ORC phosphorylation inhibits stable 1^st^ Mcm2-7 ring closing

Previous studies connected Cdt1 release to stable Mcm2-7 ring-closing (Fig. 1A, step 4) and to long-lived association of Mcm2-7 with the DNA (Ticau et al., 2017). Our finding that Mcm2-7 was less stably DNA-bound after Cdt1 release in experiments with phosORC (Fig. 5E) suggests that ORC phosphorylation decouples Cdt1 release from stable Mcm2-7 ring-closing. To investigate this possibility, we monitored closing of the Mcm2-5 gate using an Mcm2-7 complex modified with FRET donor and acceptor fluorophores on the Mcm2 and Mcm5 subunits, respectively (Figure 6A). This Mcm2-7 labeling scheme detects the open (low-FRET) and closed (high-FRET) states of the Mcm2-5 gate (Ticau et al., 2017). As previously observed (Ticau et al., 2017), individual DNA molecules in reactions containing unphosORC show effective FRET efficiency (*E*_FRET_) transitioning from the open (*E*_FRET_ < 0.30) to closed (*E*_FRET_ > 0.30) state 25-45 seconds after first Mcm2-7 arrival (Fig. 6B,C and Supp. Fig. 4A). Importantly, once the Mcm2-7 ring closes in the presence of unphosORC it predominantly remains in that state (Fig. 6C).

**Figure 6.**
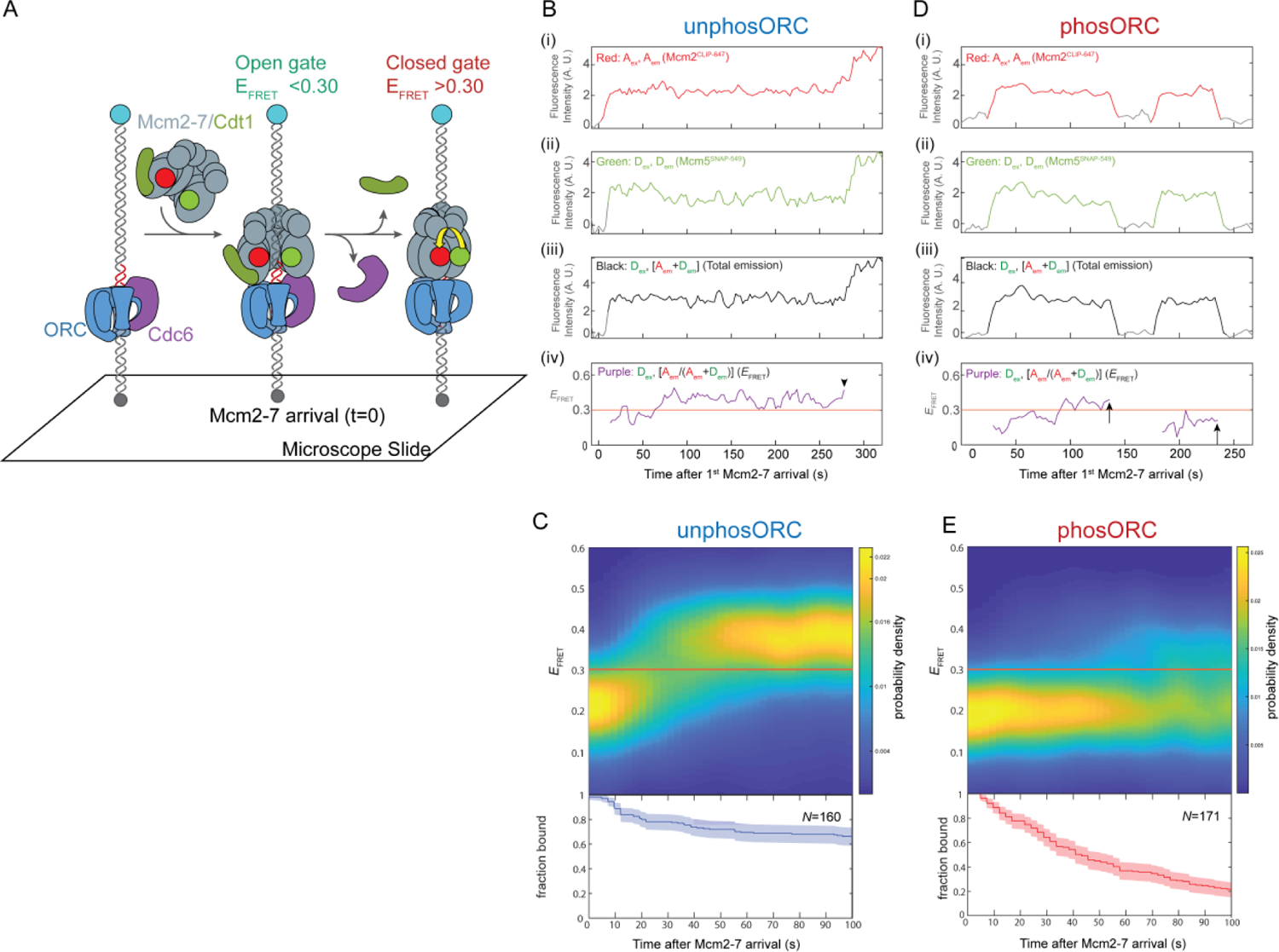
ORC phosphorylation prevents helicase ring-closing. **A.** Schematic of the Mcm2-5 ring-closing FRET assay. Mcm2-7^25FRET^ is shown in the open and closed-ring states. Mcm2 and Mcm5 were labeled with acceptor (A; red circles) and donor (D; green circles) fluorophores, respectively. Mcm2-Mcm5 ring-closure increases the proximity of the fluorophores and the FRET efficiency (*E*_FRET_). **B.** Example record of Mcm2-7^25FRET^ association with a single DNA molecule in the presence of unphosORC. Panels: Acceptor-excited acceptor emission (i, red, *A*_ex_, *A*_em_), donor-excited donor emission (ii, green, *D*_ex_, *D*_em_), donor-excited total emission (iii, black, *D*_ex_, (*A*_em_ + *D*_em_)) and calculated effective FRET efficiency (*E*_FRET_) (iv, purple, *D*_ex_, [*A*_em_ / (*A*_em_ + *D*_em_)]). AU, arbitrary units. A red line is drawn at *E*_FRET_ = 0.30, a threshold *E*_FRET_ value above which molecules are considered to be in the closed state, and below which Mcm2-7 molecules are considered to be in the open state. *E*_FRET_ is shown only during intervals when fluorescence from both labeled subunits was present, or until a 2^nd^ Mcm2-7 arrived (marked with arrowhead). Additional example records are shown in Supp. Fig. 4A. **C.** Heat map of *E*_FRET_ values vs. time after first Mcm2-7^25FRET^ binding for *N* = 160 DNA-bound 1^st^ Mcm2-7 molecules in a reaction using unphosORC (top). The plot is a two-dimensional Gaussian kernel histogram with bandwidths 5 s and 0.05 on the time and *E*_FRET_ axes, respectively, with probability densities at each time normalized independently. Only data with fluorescence from an individual double-labeled Mcm2-7 complex were included; the fraction of such complexes remaining at each time point is plotted (bottom). **D.** Example record of Mcm2-7^25FRET^ association with a single DNA molecule in the presence of phosORC. Panels are as described in (B). 1^st^ Mcm2-7 release events are marked with arrow. Additional traces are shown in Supp. Fig. 4B. **E.** Heat map of *E*_FRET_ values vs. time after first Mcm2-7^25FRET^ binding for *N* = 160 DNA-bound 1^st^ Mcm2-7 molecules in a reaction using unphosORC (top) (*N*=171). The fraction of Mcm2-7^25FRET^ complexes remaining at each time point is plotted (bottom). Plotted as described in (C).

ORC phosphorylation results in a clear defect in Mcm2-5 gate closing. Fluorescence traces from individual DNA molecules show that less than half (62/171, 36%) of the Mcm2-7 complexes recruited by phosORC achieve the closed-ring, high-*E*_FRET_ state (exhibited a minimum of two consecutive time points with *E*_FRET_ > 0.3). In contrast, with unphosORC 70% (112/160) of the 1^st^ Mcm2-7 molecules reach this state. Unlike the situation with unphosORC, 1^st^ Mcm2-7s recruited by phosORC that reach the high-*E*_FRET_ do not remain stably bound to DNA in this state. Instead, we see that Mcm2-7 durations in the closed-ring state with phosORC are short-lived (median = 17s) and either return to the open-ring, low-*E*_FRET_ state or are released from the DNA directly (presumably with a brief return to the open state to allow DNA to exit that is too short to detect). Analysis of aggregated *E*_FRET_ data from many 1^st^ Mcm2-7 molecules recruited by either phosORC or unphosORC further illustrates their different fates. In contrast to the predominant transition of the Mcm2-7s from the open to the closed state observed with unphosORC, when ORC is phosphorylated Mcm2-7s largely remain in the open state (Fig. 6, compare C and E; Supp. Fig. 5A and B). Consistent with a lack of a stably closed state, when ORC is phosphorylated, we observe a comparatively rapid loss of recruited Mcm2-7s (Fig. 6E, lower panel). In contrast, relatively fewer Mcm2-7 molecules are released after the transition to the high-*E*_FRET_ state in unphosORC reactions (Fig. 6C, lower panel). Together, these data show that Mcm2-7 recruited by phosORC is not prevented from attaining the closed-ring state but, once attained, that state is unstable resulting in the more rapid release of these molecules from the DNA.

To assess whether the same two *E*_FRET_ states were accessed by Mcm2-7 regardless of ORC phosphorylation state, we plotted the Mcm2-7 *E*_FRET_ values for various time windows (0-15, 15-50, and 50-100 s after Mcm2-7 arrival) for experiments with unphosORC and phosORC (Supp Fig. 5). Although the fraction of molecules in the low- and high-*E*_FRET_ states were not the same for phosORC and unphosORC, the positions of the two main *E*_FRET_ peaks was similar across all of the experiments. To assess the similarity more rigorously, we independently fit each of the six *E*_FRET_ distributions to a two-component Gaussian mixture model (Supp. Fig. 5A and 5B). All distributions fit to the same two-state model with similar values for the *E*FRET center positions of the two states (∼0.20 and ∼0.37, Supp. Fig. 5C and 5D). This analysis is consistent with the idea that Mcm2-7 moves between similar open and closed states whether or not ORC was phosphorylated.

As with the effects of phosphorylation of individual ORC subunits on the two Cdt1-release pathways, we asked whether phosphorylation of Orc2 or Orc6 caused the ring-closing defect. We found that phosphorylation of either Orc2 or Orc6 strongly inhibits stable Mcm2-7 ring closing (Supp. Fig. 6A and 6B). Consistent with this single-molecule data, we also found either Orc2 or Orc6 phosphorylation was sufficient to inhibit helicase loading in ensemble helicase-loading assays (Supp. Fig. 6C). Thus, unlike the choice of Cdt1-release pathway which is specific to Orc6 phosphorylation, either Orc2 or Orc6 phosphorylation inhibits stable Mcm2-7 ring closing.

### Phosphorylated ORC fails to form the MO intermediate

After the first Cdt1 dissociates, ORC releases its initial interaction with the C-terminal domains of the 1^st^ Mcm2-7 and binds the N-terminal domains of Mcm2-7 to form the MO complex (Fig. 1A, steps 4-5, (Gupta et al., 2021; Miller et al., 2019). This interface primarily involves interactions between Orc6 and the N-terminal regions of the Mcm2 and Mcm5 subunits in the closed-gate conformation. Consistent with the MO complex being associated with the closed-ring state, Mcm2-7 complexes that form this intermediate have much longer half-lives on the DNA than Mcm2-7 which fail to form the MO (Gupta et al. 2021).

Given the absence of stable Mcm2-5 ring-closing when ORC is phosphorylated, and the observation that the resolved MO structure involves the closed Mcm2-5 gate (Miller et al., 2019), we asked if ORC phosphorylation inhibited MO complex formation. To assay MO complex formation in a single-molecule assay, we measured FRET from ORC modified at the C-terminus of Orc6 with a green-excited donor fluorophore (ORC^6C-DY549^) to Mcm2-7 modified at the N-terminus of Mcm3 with a red-excited acceptor fluorophore (Mcm2-7^3N-650^, Fig. 7A). These modified proteins have previously been shown to exhibit FRET upon MO complex formation (Gupta et al., 2021). When ORC was unphosphorylated, 21 ± 3% (46/223) of 1^st^ Mcm2-7 landings showed the high-*E*_FRET_ state (>0.5) associated with MO-complex formation (Fig. 7B and 7D, Supp. Fig. 5A). In contrast, when ORC was phosphorylated none (0/162) of Mcm2-7 binding events exhibited the high-*E*_FRET_ state (Fig. 7C and 7D, Supp. Fig. 7B). As with ring closing, phosphorylation of either Orc2 or Orc6 strongly inhibited MO-complex formation, although Orc2 did not show complete inhibition (Fig. 7D).

**Fig 7.**
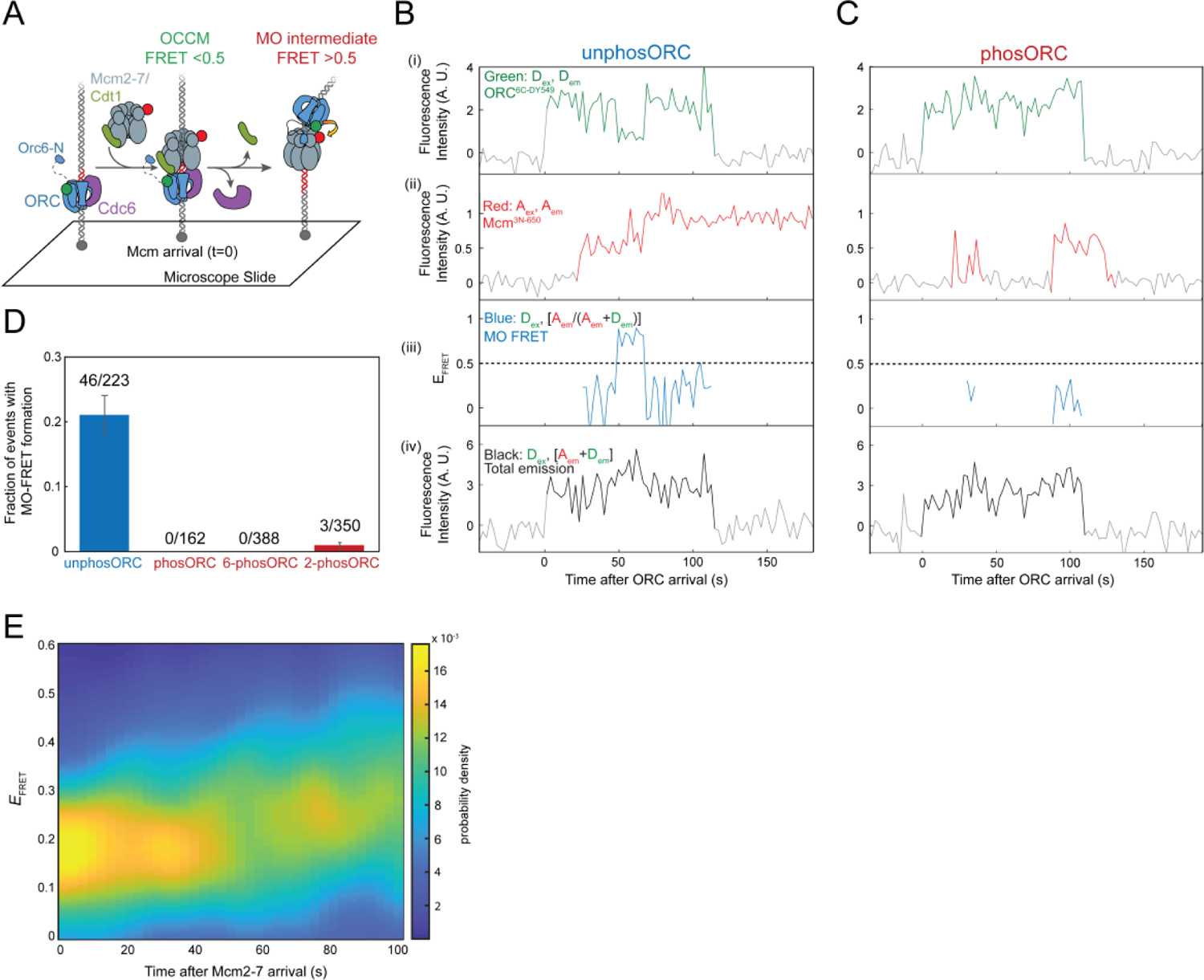
ORC phosphorylation prevents MO formation which is required for ring closing. **A.** Schematic of experiment to monitor MO-complex formation by FRET. The C-terminus of Orc6 is labeled using a green donor fluorophore (ORC^6C-549^) and the N-terminus of Mcm3 is labeled using a red acceptor fluorophore (Mcm2-7^3N-650^). When ORC and Mcm2-7 are in the OM state (e.g. in the OCCM), the dyes are far apart and *E*_FRET_ is low. Upon MO complex formation, the donor and acceptor fluorophores are in close proximity and *E*_FRET_ is high. **B.** Example record of a single-molecule helicase loading assay with Mcm2-7^3N-650^ and unphosORC^6C-549^ with a single DNA molecule. Panels: Acceptor-excited acceptor emission (i, red, *A*_ex_, *A*_em_), donor-excited donor emission (ii, green, *D*_ex_, *D*_em_), donor-excited total emission (iii, black, *D*_ex_, (*A*_em_ + *D*_em_)) and calculated effective FRET efficiency (*E*_FRET_) (iv, purple, *D*_ex_, [*A*_em_ /(*A*_em_ + *D*_em_)], only shown when Mcm2-7^3N-650^ and unphosORC^6C-549^ are both present). AU, arbitrary units. A dashed line is drawn at *E*_FRET_ = 0.5, a threshold used to define low *E*_FRET_ (*E*_FRET_ < 0.5, OM) and high *E*_FRET_ (*E*_FRET_ ≥ 0.5, MO) states. This record is representative of the 46/223 Mcm2-7 landings that showed MO *E*_FRET_ with unphosORC. Additional example records are shown in Supp. Fig. 7A. **C.** Same as (B), but with phosORC^6C-549^. Additional example records are shown in Supp. Fig. 7B. **D.** The percentage (± SEM) of Mcm2-7-DNA landings that show MO-FRET (i.e., *E*_FRET_ ≥ 0.5 for 2+ frames) for wild-type ORC^6C-549^ without CDK treatment (blue), or CDK-treated (red) wild-type ORC^6C-549^ or mutant ORC constructs that cannot be phosphorylated on Orc2 (6-phosORC^6C-549^) or Orc6 (2-phosORC^6C-549^). **E.** Heat map of *E*_FRET_ vs time in the ring-closing assay (as illustrated in Fig. 6A) for the unphosphorylated ORC^6N-Δ119^ mutant. Results plotted as in Fig. 6C (N=100).

If formation of the MO complex is required for stable Mcm2-7 ring closing, then other mutants that inhibit MO formation should have the same impact on ring closing as ORC phosphorylation. To test this hypothesis, we incorporated an ORC with a mutation that removed the N-terminal 119 amino acids of Orc6 (ORC^6N-Δ119^) into the Mcm2-7 ring-closing assay. Prior studies showed that this deleting this region of Orc6 inhibits MO complex formation (Gupta et al., 2021; Miller et al., 2019). Consistent with our hypothesis, we found that, unphosORC^6N-Δ119^ resulted in similar strong inhibition of stable Mcm2-7 ring closing (Fig. 7E) as we observe with phosORC (Fig. 6C, right). Based on our findings, we conclude that MO-complex formation is required for stable closing of the first Mcm2-7 ring.

## Discussion

The studies presented here identify multiple steps of helicase loading that are impacted by ORC phosphorylation. When ORC is phosphorylated, OCCM formation (Fig. 1A, step 2) is not blocked, but it is slowed. More significantly, ORC phosphorylation fully inhibits the helicase loading pathway between recruitment of the first and second Mcm2-7 complexes (Fig. 1A, steps 3-7). During this interval, ORC phosphorylation increases the fraction of Mcm2-7/Cdt1 recruitment events that terminate by the non-productive simultaneous release of Mcm2-7 and Cdt1 (Fig. 5A, left). Importantly, the phosORC-recruited Mcm2-7 complexes that remain DNA-associated after Cdt1 release do not reach a stable closed-ring state or form the MO complex (Fig. 1A, steps 4 and 5), resulting in their release prior to second Mcm2-7 recruitment. Our observations have implications both for how ORC phosphorylation blocks helicase loading and for the mechanism of uninhibited loading.

### ORC phosphorylation prevents stable closing of the first Mcm2-7 ring by inhibiting MO formation

Our studies identify stable closing of the first Mcm2-7 ring around the origin DNA as the key failure point in helicase loading when ORC is phosphorylated (Figs. 6 and 7). Previous studies have shown that Mcm2-7 is held in the open state while Cdt1 is bound (Frigola et al., 2017; Sun et al., 2013; Ticau et al., 2017; Yuan et al., 2017). In reactions with unphosORC, closing of the Mcm2-7 ring around DNA is kinetically similar to release of Cdt1 (Ticau et al. 2017). These data suggested that Cdt1 release is sufficient for stable Mcm2-7 ring closing. In contrast to this model, after Cdt1 dissociates from 1^st^ Mcm2-7 during reactions using phosORC, retained Mcm2-7 rings never reach a stably-closed state. Instead, we observe that Mcm2-7 is either released before or shortly after reaching the closed state. In the latter case, we assume that during release the helicase passes through the open state but for too short a time for this to be detected. This conclusion is supported both by examination of individual complexes (Fig. 6D and Supp. Fig. 4) and by population analysis showing representation of the open and closed FRET states ≥ 50s after Mcm2-7 arrival (Fig. 6E; Supp. Fig. 5B). Thus, in the presence of phosORC (and likely also unphosORC, as discussed below), Cdt1 release is not sufficient to achieve stable Mcm2-7 ring closing.

Based on our findings, we propose that MO complex formation, which follows Cdt1 release, is required to stabilize the closed-ring state of the first Mcm2-7 complex (Fig. 8). Consistent with this hypothesis, structural studies show that Orc6 binds across the closed Mcm2-5 gate in the MO complex (Miller et al. 2019). Further, when MO complex formation is inhibited, either by ORC phosphorylation or deletion of the Orc6 N-terminal domain, stable closing of the Mcm2-5 gate is not observed (Figs. 6 and 7E). Our hypothesis that MO complex formation is required for stable Mcm2-7 ring closing is also consistent with the previously determined kinetics of helicase loading: first, MO formation and Mcm2-7 ring closing occur with similar timing with unmodified ORC; and second, interfering with MO formation (with ORC^6N-Δ119^) reduces the dwell times of first Mcm2-7 complexes on the DNA (Gupta et. al., 2021). Finally, the lack of MO complex formation also explains the lack of second Cdc6 and Cdt1-Mcm2-7 recruitment events when ORC is phosphorylated, as MO complex is required for both events (Fig. 3, Gupta et al., 2021; Miller et al., 2019).

**Figure 8.**
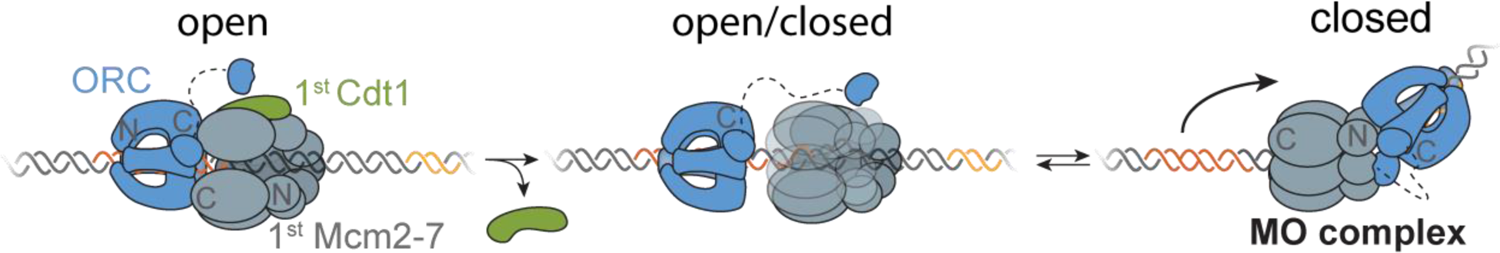
A two-step model for closing of the first Mcm2-7 ring. We propose that Cdt1 release allows Mcm2-7 to access both the open and closed states but there is no intrinsic means for the Mcm2-7 ring to stably close. Instead, we propose that formation of the MO complex is required to stabilizes the closed state.

How might MO formation be inhibited by ORC phosphorylation? It is possible that phosphorylation interferes with subunit contacts required for MO formation. Consistent with this model, the Orc2 and Orc6 subunits are at or close to the ORC-Mcm2-7 interface in the MO complex. We note, however, that the sites of Orc2 and Orc6 CDK phosphorylation are not resolved in the MO structure (presumably because they are unstructured regions, Miller et al., 2019), making it difficult to determine whether phosphorylation directly interferes with the ORC-Mcm2-7 contacts. Alternatively, it is possible that phosphorylation leads to novel interactions that are incompatible with MO complex formation. This possibility is supported by recent structural studies showing that the N-terminal domain of phosphorylated Orc6 forms a new interaction with Orc1 and Cdc6 (Schmidt et al., 2022). We note, however, that this specific interaction would not be relevant to MO formation because Cdc6 is released from the DNA well before the time of MO formation (Gupta et al., 2021). Nevertheless, it is possible that there are other novel interactions that occur when ORC is phosphorylated that interfere with forming the MO complex.

### ORC phosphorylation decreases the rate of OCCM formation and the frequency of productive Mcm2-7-Cdt1 recruitment events

Although ORC phosphorylation inhibits stable Mcm2-7 ring closing and MO complex formation completely, we also observed phosphorylation-dependent reduction in the rate of OCCM formation (Fig. 2) and decrease in productive Cdt1-release events (Fig. 5). The phosphorylation-dependent formation of a new complex between phosOrc6 and the Orc1-Cdc6 interface (Schmidt et al., 2022) is a strong candidate to influence both events. This interaction interferes with Mcm7 binding to the ORC-Cdc6 bound to DNA and could reduce the rate of successful Mcm2-7-Cdt1 recruitment to form the OCCM. Similarly, once the OCCM is formed, it is possible that residual interactions of phosOrc6 with this region could increase the fraction of Mcm2-7-Cdt1 recruitment events in which both proteins are released simultaneously. Given the continued formation of OCCM in the presence of phosORC, the interaction of phosOrc6 with the Orc1-Cdc6 interface is likely to be either transient or frequently outcompeted by Mcm2-7-Cdt1. In addition to this phosphorylation-dependent interaction, CDK phosphorylation also inhibits an interaction between Orc6 and Cdt1 (Chen and Bell, 2011). This interaction has been observed by direct association assays (Chen and Bell, 2011) and cross-linking studies in the context of the OCCM (Yuan et al., 2017). Interference with this interaction could influence the type of Cdt1 release events that occur, and thereby altering the fraction of Mcm2-7-Cdt1 recruitment events that follow the productive pathway.

Although neither the reduced rate of OCCM formation or altered fraction of productive Cdt1 release events completely prevents helicase loading, their effects could be significant *in vitro* and *in vivo*. Although we did not detect any MO formation in our *in vitro* studies, by reducing the number of stably associated 1^st^ Mcm2-7 complexes, these effects would further reduce the potential opportunities for MO formation. *In vivo*, slower OCCM formation could enhance inhibition of helicase loading at the beginning of S-phase by giving phosphorylated Cdc6 and Mcm2-7 more time to be degraded (Cdc6) or removed from the nucleus (Mcm2-7). Similarly, increasing the fraction of non-productive Mcm2-7-Cdt1 recruitment events would provide additional opportunities for CDK-phosphorylation-dependent Mcm2-7 nuclear export (only non-DNA-bound Mcm2-7 is subject to this regulation).

### Orc2 and Orc6 phosphorylation impact helicase loading differently

Phosphorylation of Orc2 and Orc6 do not have the same impact on helicase loading. Only Orc6 phosphorylation independently alters the fraction of OCCM complexes that take the non-productive Cdt1-release pathway (Fig. 5D). Although eliminating Orc2 phosphorylation on its own has no effect on this pathway choice, elimination of both Orc2 and Orc6 phosphorylation has a stronger effect than eliminating Orc6 phosphorylation alone. This finding suggests that Orc2 phosphorylation enhances the impact of Orc6 phosphorylation on the pathway of Cdt1 release. In contrast, we observe that phosphorylation of Orc2 or Orc6 alone results in strong effects on Mcm2-7 ring closing and MO formation that are similar to that seen with fully phosphorylated ORC (Fig. 7D and Supp. Fig. 6). Consistent with this observation, *in vivo* studies show that elimination of both Orc2 and Orc6 phosphorylation sites is necessary to observe substantial re-replication of the genome (Chen and Bell, 2011).

### Implications for Mcm2-7 ring closing

In addition to providing insights into the control of helicase loading by ORC phosphorylation, our studies have important implications for the mechanism of uninhibited helicase loading. Our finding that Cdt1 release does not intrinsically lead to Mcm2-7 ring closing when ORC is phosphorylated suggests that the same is likely to be true in the absence of phosphorylation. We propose that stable closing of the first Mcm2-7 is a two-step process (Fig. 8). First, release of Cdt1 (stimulated by Mcm2-7 ATP hydrolysis, Ticau et al., 2017) allows Mcm2-7 to access the closed ring state. Our findings indicate that there is no Mcm2-7-intrinsic ‘clasp’ that holds the ring closed. Instead, Mcm2-7 accesses both the open and closed states after Cdt1 release (Fig. 6 and Supp. Fig. 5). The second step in stable closing of the first Mcm2-7 ring is MO complex formation which maintains the first Mcm2-7 in a closed state.

Our model further suggests that anytime an Mcm2-7 ring is stably closed, it requires other proteins to hold it in that state. This leads to that question of what performs this function after the MO complex dissolves. The MO complex is present when second Mcm2-7 arrives but is lost shortly thereafter. The time of MO loss correlates with the time of initial formation of double-hexamer interactions (Gupta et al., 2021). We propose that the MO complex holds the first Mcm2-7 ring closed until the second Mcm2-7 arrives at which point interactions between the two Mcm2-7 complexes keep the first Mcm2-7 ring closed. Because the first Mcm2-7 ring does not reopen at any time after its initial closure (Ticau et al., 2017), it is likely that interactions between the two Mcm2-7s stabilize the closed state of the first Mcm2-7 and displace the MO interactions. Notably, the interactions between the Mcm2-7s after MO displacement do not immediately lead to closing of the second Mcm2-7 ring (Fig. 1A, steps 8-10). This event only occurs when the Cdt1 associated with the second Mcm2-7 is released (Ticau et al., 2017). Nevertheless, we propose that Cdt1 release allows the second Mcm2-7 to access the closed-ring state and reciprocal interactions between the first Mcm2-7 and the gate of the second Mcm2-7 stabilize the closed state.

Such a model has the appeal that the MO complex would only dissolve once the second Mcm2-7 is present to maintain the closed state. Then the first Mcm2-7 ring would rapidly stabilize the closed ring of the second Mcm2-7 as soon as the release of Cdt1 allowed this state to be accessed. Consistent with the idea that the Mcm2-7 ring will not remain closed on its own, other proteins are observed to hold the Mcm2-7 ring closed at later stages of replication. Cdc45 and GINS form a bridge that holds the Mcm2-7 gate closed in the context of the active CMG helicase (Costa et al., 2014). This raises the interesting possibility that separation of the two helicases in the double hexamer during helicase activation allows reopening of the Mcm2-7 rings to allow ssDNA extrusion from the ring before Cdc45-GINS binding captures the closed-ring state. Further studies examining the relative time of separation of the two helicases, ssDNA extrusion, and Cdc45-GINS binding to the Mcm2-5 gate will be required to test this hypothesis.

## Material and Methods

### Protein purification and fluorescent labeling

Wild-type ORC was purified as previously described from the yeast strain ySDORC that expresses codon-optimized versions of the six ORC genes (Frigola et al., 2013). Unlabeled ORC with alanine substitutions at Orc2 (*orc2-6A*, alanine substitutions for S or T) or Orc6 (*orc6-4A)* CDK phosphorylation sites (Chen and Bell, 2011) were purified from the following strains: 6-phosORC, yAA01; 2-phosORC, yAA02; and no-phosORC, yAA03. Fluorescently-labeled ORC was purified from the following strains: ORC^6C-549^, ySG039; 6-phosORC^6C-549^, ySG060; and 2-phosORC^6C-549^, ySG061. Wild-type unlabeled Cdc6 was purified from BL21-DE3-Rosetta bacteria transformed with the plasmid pSKM033 and Cdc6^SORT-549^ was purified from the same bacteria transformed with pET-GSS-Cdc6. Mcm2-7 was purified from one of three strains depending on how it was fluorescently labeled: Mcm2-7^4SNAPDY649^/Cdt1^SORT549^ and Mcm2-7^4SNAP-DY649^-Cdt1 were purified from yST166; Mcm2-7^4SORTDY649^ and Mcm2-7^4SORTJF646^ were purified from yST180, Mcm2-7^25FRET^ (with Mcm2^CLIP^ and Mcm5^SNAP^, which form a FRET pair at the Mcm2-5 gate when donor/acceptor FRET dyes are used) was purified from yST229; and Mcm2-7^3N-650^ was purified from ySG024. Cdt1^SORT^ without Mcm2-7 was purified from yST103. Clb5-Cdk1 was purified from ySK119 and Sic1 was purified from BL21-DE3-Rosetta bacteria transformed with the plasmid pGEX-Sic1 (Heller et al., 2011). The yeast strains used, and their genotypes are listed in Supp. Table 2 and expression plasmids used are listed in Supp. Table 3.

Unlabeled ORC mutants were purified as described in (Tsakraklides and Bell, 2010) with the following modifications: following the Flag-antibody-coupled column (Sigma), the elutions containing ORC (typically 2-5) were pooled and incubated with 40 µl of equilibrated Anti-V5-antibody-coupled Agarose Affinity Gel (Sigma) for 30 min at 4°C with rotation, then the Anti-V5 Agarose beads were removed by spinning and pelleting according to manufacturer’s instructions. The resulting supernatant was transferred to a new tube and incubated with another 40 µl of equilibrated Anti-V5 Agarose overnight (12h) at 4°C with rotation. The Anti-V5 Agarose was removed as before and ORC purification was continued using a SPHP column (Cytiva). ORC mutants were purified as described in (Gupta et al., 2021) with the same modifications described above following the calmodulin-binding peptide affinity column.

Clb5-Cdk1 was purified as described in (Looke et al., 2017). Cdc6 and Cdc6^SORT-549^ were purified as described (Mehanna and Diffley, 2012; Ticau et al., 2015). Sic1 was purified as described (Heller et al., 2011) but with 1 mM IPTG for induction, 200 mM KCl in all buffers, and 20 mM Glutathione (pH 8) for elution.

Fluorescently labeled Mcm2-7^4SORTDY649^/Cdt1 (Fig. 4), Mcm2-7^4SORTJF646^/Cdt1^SORT549^ (Fig. 5), Cdc6^SORTDY549^, and Cdt1^SORTDY549^ (Fig. 5) were purified and labeled using an N-terminal sortase reaction as previously described (Ticau et al., 2015). Mcm2-7^25FRET^ was purified and labeled as previously described (Ticau et al., 2017). Fluorescently-labeled ORC^6C-549^, 6-phosORC^6C-549^, and 2-phosORC^6C-549^ were purified and labeled using a C-terminal sortase reaction as described in (Gupta et al., 2021). Bovine serum albumin was added to a final concentration of 1 mg/ml to all fluorescent proteins before aliquoting and flash freezing to improve long-term stability.

### *In vitro* phosphorylation of ORC

1 mM ATP and 1 mM MgOAc was added to ORC protein aliquots and then each aliquot was then divided in half. Purified Clb5-Cdk1 CDK was added to a concentration of 400 nM to one of the two tubes. Both tubes were incubated at room temperature for 20 minutes. After 20 min, purified GST-Sic1, a potent B-type CDK inhibitor in budding yeast (Schwob et al., 1994), was added to a concentration of 800 nM to both samples and incubated at room temperature for 5 min to inhibit Clb5-Cdk1 and prevent it from phosphorylating Mcm2-7 and Cdc6 when ORC is added to the full reaction. The two ORC samples were then placed on ice until they were added to the reaction.

### Western blot confirmation of ORC phospho-site mutants

Both unphosphorylated and phosphorylated (as above) samples of each purified ORC mutant protein were run on an 8% polyacrylamide gel. Phosphorylation dependent band-shifts were then visualized by western blot using custom mouse anti-Orc2 and anti-Orc6 monoclonal antibodies (anti-ORC2, SB46; anti-ORC6, SB49)

### Single-molecule fluorescence microscopy

The micromirror total internal reflection microscope used for these studies (excitation wavelengths 488, 532, and 633 nm; autofocus wavelength 785 nM) is described in (Friedman et al., 2006; Friedman and Gelles, 2015). Single molecule helicase loading reactions were performed as previously described in Ticau et al., 2015 but with the addition of triplet state quenchers to further minimize photobleaching (Friedman et al., 2006; Hoskins et al., 2011). The high-salt wash at the end of experiments was left in the slide chamber for 1 min and was then followed by a low salt wash consisting of reaction buffer, triplet state quenchers and oxygen scavengers before imaging the remaining loaded helicases. Slides and coverslips were prepared as previously described (Ticau et al., 2015) with the following changes. The two types of PEG were applied at different concentrations: mPEG-silane-2000 (from Creative PEGWorks) was used at a 50 mg/mL concentration, and biotin-mPEG-silane-3400 (Laysan Bio) was used at 0.1 mg/ml (500:1 weight ratio). Before flowing streptavidin and DNA into the chamber (to attached DNA to the surface), streptavidin-coated broadly-fluorescent beads (0.04 µm, Invitrogen TransFluoSpheres) were added (at a dilution of 1:200,000-1:400,000 into 10mM Tris-HCL, pH 8, 0.5mM BSA, 1% NP-40) and incubated until there were 2-5 beads per 65 µM diameter field of view. These beads served as fiducial markers for thermal drift correction during image analysis. We identified DNA molecule locations (before addition of helicase-loading proteins) by acquiring 5 images with a 488 nm excitation, and fluorescent impurity locations by acquiring 10-30 images with simultaneous 532 nm and 633 nm excitation). Any “DNA” locations that co-localized with impurity fluorescence were omitted from analysis.

Helicase loading reactions with labeled Mcm2-7, labeled Cdt1, labeled and unlabeled ORC, and unlabeled Cdc6 or doubly-labeled Mcm2-7^25FRET^ contained 2.5 nM ORC, 10 nM Cdc6, and 2.5 nM Mcm2-7-Cdt1. Fluorescent Cdc6 was spun at 200,000 × *g* for 20 min before use to pellet aggregates.

### Mcm2-5 gate FRET assays

FRET experiments were performed as described (Ticau et al., 2017) but with the following modification: After addition of the helicase loading proteins to the chamber, we initially acquired sequential 1 s duration images of 488 nm, 532 nm, and 633 nm excitation repeated 4 - 10 times. For the remainder of the reaction, image acquisition was switched to alternating between green and red excitation for 600 frames of each color (including a ∼0.3 s time interval between frames). Reactions were typically monitored for ∼26 min. The initial 4-10 green frames were then prepended to the subsequent 600 green frames, and the same was done for initial and subsequent red-excited frames.

### Single-molecule data analysis

Single-molecule videos were converted into numerical tables of fluorescent protein-DNA colocalization times using custom software and algorithms for automatic spot detection as described previously (Friedman and Gelles, 2015; Ticau et al., 2015) and available here: https://github.com/gelles-brandeis/CoSMoS_Analysis). The automatic spot detection results were checked, and manually corrected when necessary. The resulting tables of protein-DNA binding events were used to determine time-to-first binding kinetics (described below), the survival curves shown in Fig. 4, 5, 6, and the Cdt1-release fraction bar graph in Fig. 6, and to select the intervals where both dyes in the FRET-pair were present in Fig. 6, (codes provided link/site). Identification of 2^nd^ Mcm2-7 binding (Fig. 3, Fig 6) was done manually.

### Time-to-first-binding analysis

Mcm2-7 time to first binding curves were plotted and fit to a model that is the sum of exponential specific binding and exponential non-specific background binding as described in (Friedman and Gelles, 2015). The rate of background binding was determined by monitoring Mcm2-7 binding at locations that did not contain a DNA molecule and were >5 pixels away from any DNA molecule (1000 such non-DNA locations evenly selected throughout the field of view were evaluated). Fitting the time to first binding data at the non-DNA locations yielded an association rate constant for non-specific binding of Mcm2-7 to the surface of the slide, *k_n,s_*. Fitting the Mcm2-7 binding at DNA locations yielded the apparent first-order association rate constant, *k_a_*, and the fraction of DNAs that were active (i.e., capable of binding Mcm2-7), *A_f_*.

### Mcm2-7 ring-closing FRET data analysis

Mcm2-7^25FRET^ experiments were analyzed as described previously (Ticau et al 2017) with the following modifications. Donor and acceptor fluorescence intensity recorded during donor excitation were background corrected by sampling and smoothing the intensity of the area surrounding around each DNA. Automated spot detection was used to determine time-intervals on individual DNA molecules where an Mcm2-7 containing both Mcm2-donor (DY549) and Mcm5-acceptor (Dy649) was present; the first and last frame from each spot were discarded. These intervals were then manually checked for 2^nd^ Mcm2-7 arrival and the apparent E_FRET_ was calculated only during the period before the 2^nd^ Mcm2-7 arrived. For scoring ring-closing, an Mcm2-7 ring was considered closed if the E_FRET_ signal was >0.3 for 2 or more frames.

### Two-dimensional *E*_FRET_ heat maps

“The arrival of each 1^st^ Mcm2-7^25FRET^ molecule was set as time zero. All events were then plotted as a two-dimensional Gaussian kernel histogram with bandwidths of 5 s on the time axis and 0.05 on the *E*_FRET_ axes, and were normalized so that the probability density in each 2.7 s time slice integrated to one” (Champasa et al., 2019; Ticau et al., 2017).

### Fitting one-dimensional histograms of *E*_FRET_ data

The *E*_FRET_ data for Mcm2-7 ring-closing in the presence of unphosphorylated or phosphorylated ORC was selected from three time intervals after 1^st^ Mcm2-7 arrival: 0 – 15 s, 15 – 50 s, and 50-100 s. *E*_FRET_ values from 0.1-0.7 (which constitute >99% of observations, for unphosORC, and >97% for phosORC) from each interval were independently fit to the two-component Gaussian mixture probability density function

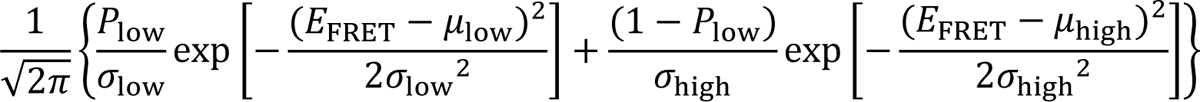

where *P*_low_ is the fractional amplitude of the low *E*_FRET_ component, µ_low_ < µ_high_ are the mean *E*_FRET_ values of the low and high components, and σ_low_, σ_high_ are the standard deviations of the low and high *E*_FRET_ components. Standard errors of the fit parameters were computed by bootstrapping (1,000 samples).

### MO formation single-molecule FRET assays

MO FRET assays were performed as described (Gupta et al., 2021). The MO complex was considered to be formed if the calculated *E*_FRET_ was greater than 0.5 for two consecutive frames.

### Data Availability

Source data for the single-molecule experiments is provided as Matlab “intervals” files that can be read and manipulated by the program imscroll, available at https://github.com/gelles-brandeis/CoSMoS_Analysis). The source data are archived at: 10.5281/zenodo.7455772.

## Supporting information

Amasino et al. Supplemental Figures and Tables

## Acknowledgements

We are grateful to members of the Bell Laboratory for useful discussions. We thank Annie Zhang for comments on the manuscript. This work was supported by NIH grants R01 GM147960 (SPB and JG) and R01 GM81648 (JG). AA and SG were both supported in part by an NIH Pre-Doctoral Training Grant (T32 GM007287), a MathWorks Science Fellowship (SG), and a Paul and Cleo Schimmel Fellowship (AA). SPB is an investigator with the Howard Hughes Medical Institute. This work was supported in part by the Koch Institute Support Grant P30-CA14051 from the NCI. We thank the Biopolymers core of the Koch Institute Swanson Biotechnology Center for technical support.

## References

1. Aparicio OM, Weinstein DM, Bell SP. 1997. Components and dynamics of DNA replication complexes in S. cerevisiae: redistribution of MCM proteins and Cdc45p during S phase. Cell 91:59–69.

2. Arias EE, Walter JC. 2007. Strength in numbers: preventing rereplication via multiple mechanisms in eukaryotic cells. Genes & development 21:497–518. doi:10.1101/gad.1508907

3. Bell SP, Labib K. 2016. Chromosome Duplication in Saccharomyces cerevisiae. Genetics 203:1027–1067. doi:10.1534/genetics.115.186452

4. Bochman ML, Schwacha A. 2008. The Mcm2-7 complex has in vitro helicase activity. Molecular cell 31:287–293. doi:10.1016/j.molcel.2008.05.020

5. Champasa K, Blank C, Friedman LJ, Gelles J, Bell SP. 2019. A conserved Mcm4 motif is required for Mcm2-7 double-hexamer formation and origin DNA unwinding. eLife 8:1822. doi:10.7554/elife.45538

6. Chen S, Bell SP. 2011. CDK prevents Mcm2-7 helicase loading by inhibiting Cdt1 interaction with Orc6. Genes & development 25:363–372. doi:10.1101/gad.2011511

7. Costa A, Renault L, Swuec P, Petojevic T, Pesavento JJ, Ilves I, MacLellan-Gibson K, Fleck RA, Botchan MR, Berger JM. 2014. DNA binding polarity, dimerization, and ATPase ring remodeling in the CMG helicase of the eukaryotic replisome. eLife 3:e03273. doi:10.7554/elife.03273

8. Coster G, Diffley JFX. 2017. Bidirectional eukaryotic DNA replication is established by quasi-symmetrical helicase loading. *Science (New York*, NY) 357:314–318. doi:10.1126/science.aan0063

9. Donovan S, Harwood J, Drury LS, Diffley JFX. 1997. Cdc6p-dependent loading of Mcm proteins onto pre-replicative chromatin in budding yeast. Proc National Acad Sci 94:5611–5616. doi:10.1073/pnas.94.11.5611

10. Drury LS, Perkins G, Diffley JF. 2000. The cyclin-dependent kinase Cdc28p regulates distinct modes of Cdc6p proteolysis during the budding yeast cell cycle. Current biology : CB 10:231–240.

11. Drury LS, Perkins G, Diffley JF. 1997. The Cdc4/34/53 pathway targets Cdc6p for proteolysis in budding yeast. The EMBO journal 16:5966–5976. doi:10.1093/emboj/16.19.5966

12. Elsasser S, Chi Y, Yang P, Campbell JL. 1999. Phosphorylation Controls Timing of Cdc6p Destruction: A Biochemical Analysis. Mol Biol Cell 10:3263–3277. doi:10.1091/mbc.10.10.3263

13. Evrin C, Clarke P, Zech J, Lurz R, Sun J, Uhle S, Li H, Stillman B, Speck C. 2009. A double-hexameric MCM2-7 complex is loaded onto origin DNA during licensing of eukaryotic DNA replication. Proceedings of the National Academy of Sciences of the United States of America 106:20240–20245. doi:10.1073/pnas.0911500106

14. Feng X, Noguchi Y, Barbon M, Stillman B, Speck C, Li H. 2021. The structure of ORC–Cdc6 on an origin DNA reveals the mechanism of ORC activation by the replication initiator Cdc6. Nat Commun 12:3883. doi:10.1038/s41467-021-24199-1

15. Friedman LJ, Chung J, Gelles J. 2006. Viewing dynamic assembly of molecular complexes by multi-wavelength single-molecule fluorescence. Biophysical journal 91:1023–1031. doi:10.1529/biophysj.106.084004

16. Friedman LJ, Gelles J. 2015. Multi-wavelength single-molecule fluorescence analysis of transcription mechanisms. *Methods (San Diego*, Calif) 86:27–36. doi:10.1016/j.ymeth.2015.05.026

17. Frigola J, He J, Kinkelin K, Pye VE, Renault L, Douglas ME, Remus D, Cherepanov P, Costa A, Diffley JFX. 2017. Cdt1 stabilizes an open MCM ring for helicase loading. Nature communications 8:15720. doi:10.1038/ncomms15720

18. Frigola J, Remus D, Mehanna A, Diffley JFX. 2013. ATPase-dependent quality control of DNA replication origin licensing. Nature 495:339–343. doi:10.1038/nature11920

19. Gaillard H, García-Muse T, Aguilera A. 2015. Replication stress and cancer. Nat Rev Cancer 15:276–289. doi:10.1038/nrc3916

20. Gupta S, Friedman LJ, Gelles J, Bell SP. 2021. A helicase-tethered ORC flip enables bidirectional helicase loading. Elife 10. doi:10.7554/elife.74282

21. Heller RC, Kang S, Lam WM, Chen S, Chan CS, Bell SP. 2011. Eukaryotic origin-dependent DNA replication in vitro reveals sequential action of DDK and S-CDK kinases. Cell 146:80– 91. doi:10.1016/j.cell.2011.06.012

22. Hoskins AA, Friedman LJ, Gallagher SS, Crawford DJ, Anderson EG, Wombacher R, Ramirez N, Cornish VW, Gelles J, Moore MJ. 2011. Ordered and dynamic assembly of single spliceosomes. *Science (New York*, NY) 331:1289–1295. doi:10.1126/science.1198830

23. Labib K, Diffley JF, Kearsey SE. 1999. G1-phase and B-type cyclins exclude the DNA-replication factor Mcm4 from the nucleus. Nature cell biology 1:415–422. doi:10.1038/15649

24. Liku ME, Nguyen VQ, Rosales AW, Irie K, Li JJ. 2005. CDK phosphorylation of a novel NLS-NES module distributed between two subunits of the Mcm2-7 complex prevents chromosomal rereplication. Molecular biology of the cell 16:5026–5039. doi:10.1091/mbc.e05-05-0412

25. Looke M, Maloney MF, Bell SP. 2017. Mcm10 regulates DNA replication elongation by stimulating the CMG replicative helicase. Genes & development 31:291–305. doi:10.1101/gad.291336.116

26. Marahrens Y, Stillman B. 1992. A yeast chromosomal origin of DNA replication defined by multiple functional elements. *Science (New York*, NY) 255:817–823.

27. Mehanna A, Diffley JFX. 2012. Pre-replicative complex assembly with purified proteins. Methods 57:222–226. doi:10.1016/j.ymeth.2012.06.008

28. Miller TCR, Locke J, Greiwe JF, Diffley JFX, Costa A. 2019. Mechanism of head-to-head MCM double-hexamer formation revealed by cryo-EM. Nature 575:704–710. doi:10.1038/s41586-019-1768-0

29. Nguyen VQ, Co C, Irie K, Li JJ. 2000. Clb/Cdc28 kinases promote nuclear export of the replication initiator proteins Mcm2-7. Current biology : CB 10:195–205.

30. Nguyen VQ, Co C, Li JJ. 2001. Cyclin-dependent kinases prevent DNA re-replication through multiple mechanisms. Nature 411:1068–1073. doi:10.1038/35082600

31. Phizicky DV, Berchowitz LE, Bell SP. 2018. Multiple kinases inhibit origin licensing and helicase activation to ensure reductive cell division during meiosis. eLife 7:e33309. doi:10.7554/elife.33309

32. Prioleau M-N, MacAlpine DM. 2016. DNA replication origins—where do we begin? Gene Dev 30:1683–1697. doi:10.1101/gad.285114.116

33. Randell JCW, Bowers JL, Rodríguez HK, Bell SP. 2006. Sequential ATP Hydrolysis by Cdc6 and ORC Directs Loading of the Mcm2-7 Helicase. Molecular cell 21:29–39. doi:10.1016/j.molcel.2005.11.023

34. Remus D, Beuron F, Tolun G, Griffith JD, Morris EP, Diffley JFX. 2009. Concerted Loading of Mcm2–7 Double Hexamers around DNA during DNA Replication Origin Licensing. Cell 139:719–730. doi:10.1016/j.cell.2009.10.015

35. Samel SA, Fernández-Cid A, Sun J, Riera A, Tognetti S, Herrera MC, Li H, Speck C. 2014. A unique DNA entry gate serves for regulated loading of the eukaryotic replicative helicase MCM2-7 onto DNA. Genes & development 28:1653–1666. doi:10.1101/gad.242404.114

36. Schmidt JM, Yang R, Kumar A, Hunker O, Seebacher J, Bleichert F. 2022. A mechanism of origin licensing control through autoinhibition of S. cerevisiae ORC·DNA·Cdc6. Nat Commun 13:1059. doi:10.1038/s41467-022-28695-w

37. Schwob E, Böhm T, Mendenhall MD, Nasmyth K. 1994. The B-type cyclin kinase inhibitor p40SIC1 controls the G1 to S transition in S. cerevisiae. Cell 79:233–244. doi:10.1016/0092-8674(94)90193-7

38. Seki T, Diffley JF. 2000. Stepwise assembly of initiation proteins at budding yeast replication origins in vitro. Proceedings of the National Academy of Sciences of the United States of America 97:14115–14120. doi:10.1073/pnas.97.26.14115

39. Siddiqui K, On KF, Diffley JFX. 2013. Regulating DNA Replication in Eukarya. Csh Perspect Biol 5:a012930. doi:10.1101/cshperspect.a012930

40. Stillman B. 2022. The remarkable gymnastics of ORC. Elife 11:e76475. doi:10.7554/elife.76475

41. Sun J, Evrin C, Samel SA, Fernández-Cid A, Riera A, Kawakami H, Stillman B, Speck C, Li H. 2013. Cryo-EM structure of a helicase loading intermediate containing ORC-Cdc6-Cdt1-MCM2-7 bound to DNA. Nature structural & molecular biology 20:944–951. doi:10.1038/nsmb.2629

42. Tanaka S, Umemori T, Hirai K, Muramatsu S, Kamimura Y, Araki H. 2007. CDK-dependent phosphorylation of Sld2 and Sld3 initiates DNA replication in budding yeast. Nature 445:328– 332. doi:10.1038/nature05465

43. Ticau S, Friedman LJ, Champasa K, Corrêa IR, Gelles J, Bell SP. 2017. Mechanism and timing of Mcm2–7 ring closure during DNA replication origin licensing. Nat Struct Mol Biol 24:309– 315. doi:10.1038/nsmb.3375

44. Ticau S, Friedman LJ, Ivica NA, Gelles J, Bell SP. 2015. Single-molecule studies of origin licensing reveal mechanisms ensuring bidirectional helicase loading. Cell 161:513–525. doi:10.1016/j.cell.2015.03.012

45. Tsakraklides V, Bell SP. 2010. Dynamics of pre-replicative complex assembly. Journal of Biological Chemistry 285:9437–9443. doi:10.1074/jbc.m109.072504

46. Wilmes GM, Bell SP. 2002. The B2 element of the Saccharomyces cerevisiae ARS1 origin of replication requires specific sequences to facilitate pre-RC formation. Proceedings of the National Academy of Sciences of the United States of America 99:101–106. doi:10.1073/pnas.012578499

47. Yuan Z, Riera A, Bai L, Sun J, Nandi S, Spanos C, Chen ZA, Barbon M, Rappsilber J, Stillman B, Speck C, Li H. 2017. Structural basis of Mcm2-7 replicative helicase loading by ORC-Cdc6 and Cdt1. Nat Struct Mol Biol 24:316–324. doi:10.1038/nsmb.3372

48. Zegerman P, Diffley JFX. 2007. Phosphorylation of Sld2 and Sld3 by cyclin-dependent kinases promotes DNA replication in budding yeast. Nature 445:281–285. doi:10.1038/nature05432

