## Supplementary material for "Regulation of replication origin licensing by ORC phosphorylation reveals a two-step mechanism for Mcm2-7 ring closing": Amasino et al. Supplemental Figures and Tables

**Amasino et al., Supplementary Figures**

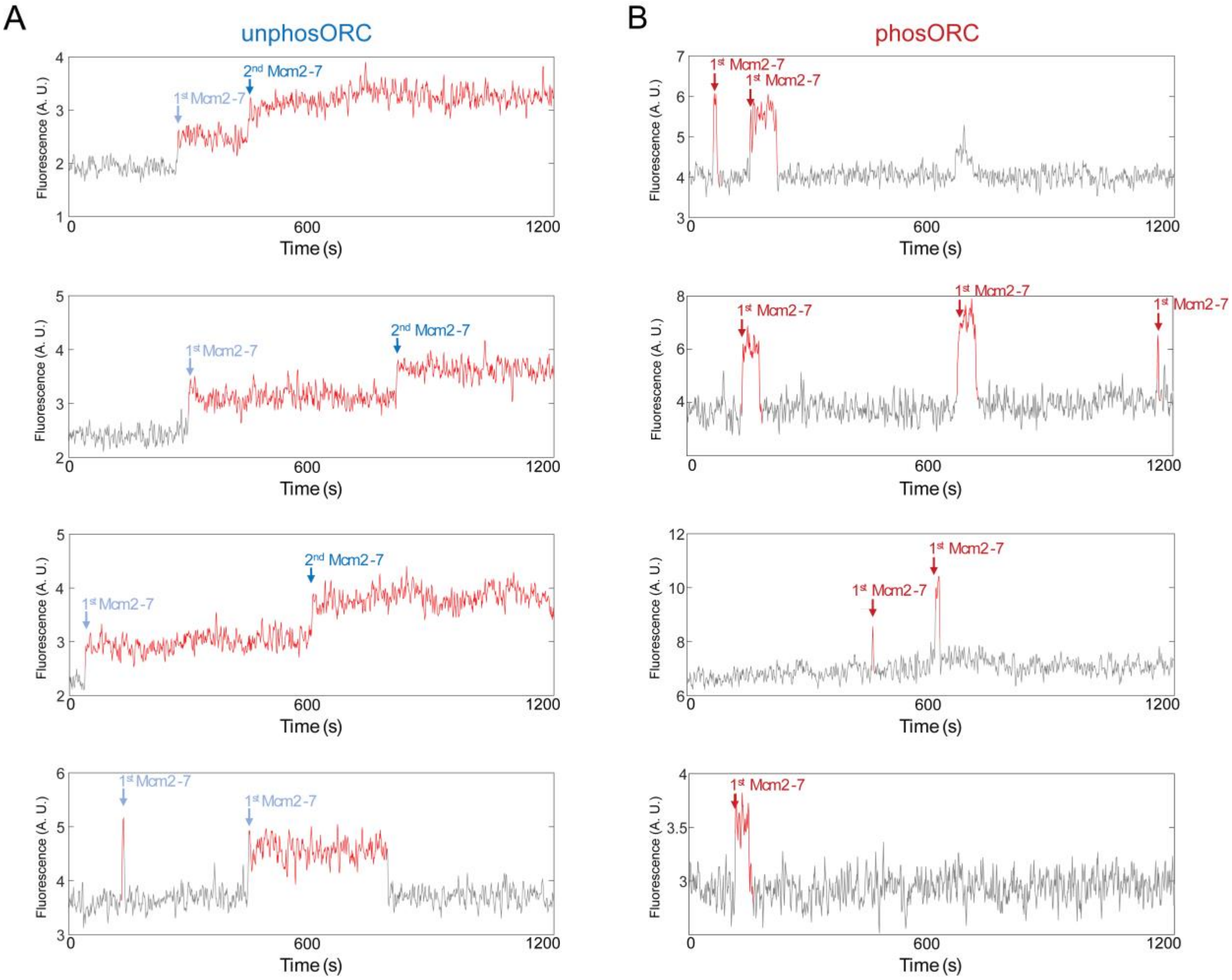


**Supplementary Figure 1. Additional fluorescence records showing Mcm2-7$^{4SNAP\text{-}JF646}$ association with single DNA molecules.**

**A.** Additional example records of Mcm2-7$^{4SNAP\text{-}JF646}$ associations with an individual DNA molecule in the presence of unphosORC (experiment same as in Fig. 1), plotted as in Fig. 2A.

**B.** Additional example records of Mcm2-7$^{4SNAP\text{-}JF646}$ associations with an individual DNA molecule in the presence of phosORC (experiment same as in Fig. 1), plotted as in Fig. 2A.

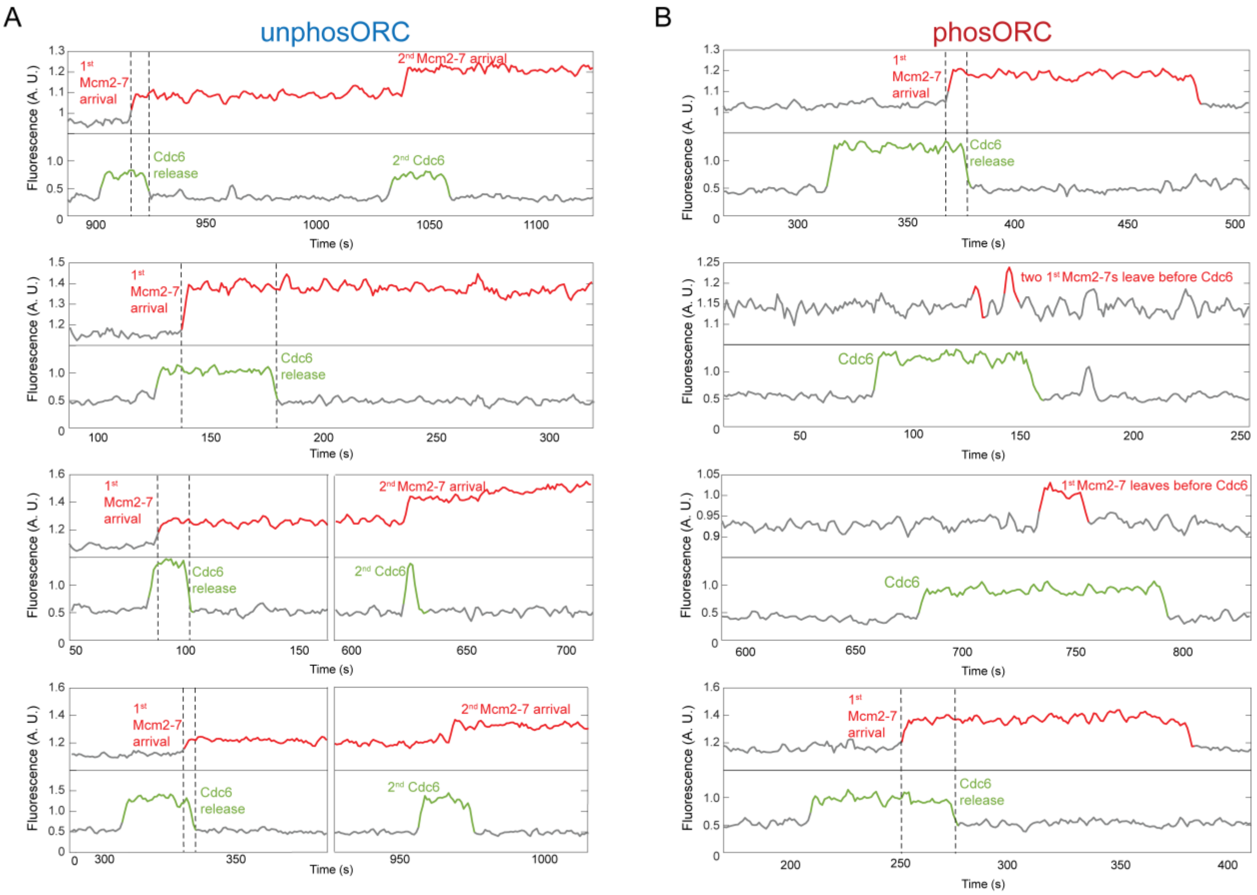


**Supplementary Figure 2. Additional fluorescence records of Mcm2-7 and Cdc6 interactions with single DNA molecules.**

A. Additional example records of Mcm2-7$^{4SNAP\text{-}DY649}$ (red) and Cdc6$^{Sort\text{-}549}$ (green) records from a single-molecule helicase-loading experiment with unlabeled Cdt1 and unphosORC. Dashed lines and plotting are as described in Fig. 4B.

B. Additional Mcm2-7$^{4SNAP\text{-}DY649}$ (red) and Cdc6$^{Sort\text{-}549}$ (green) records from a single-molecule helicase-loading experiment with phosORC. Dashed lines and plotting are as described in Fig. 4B.

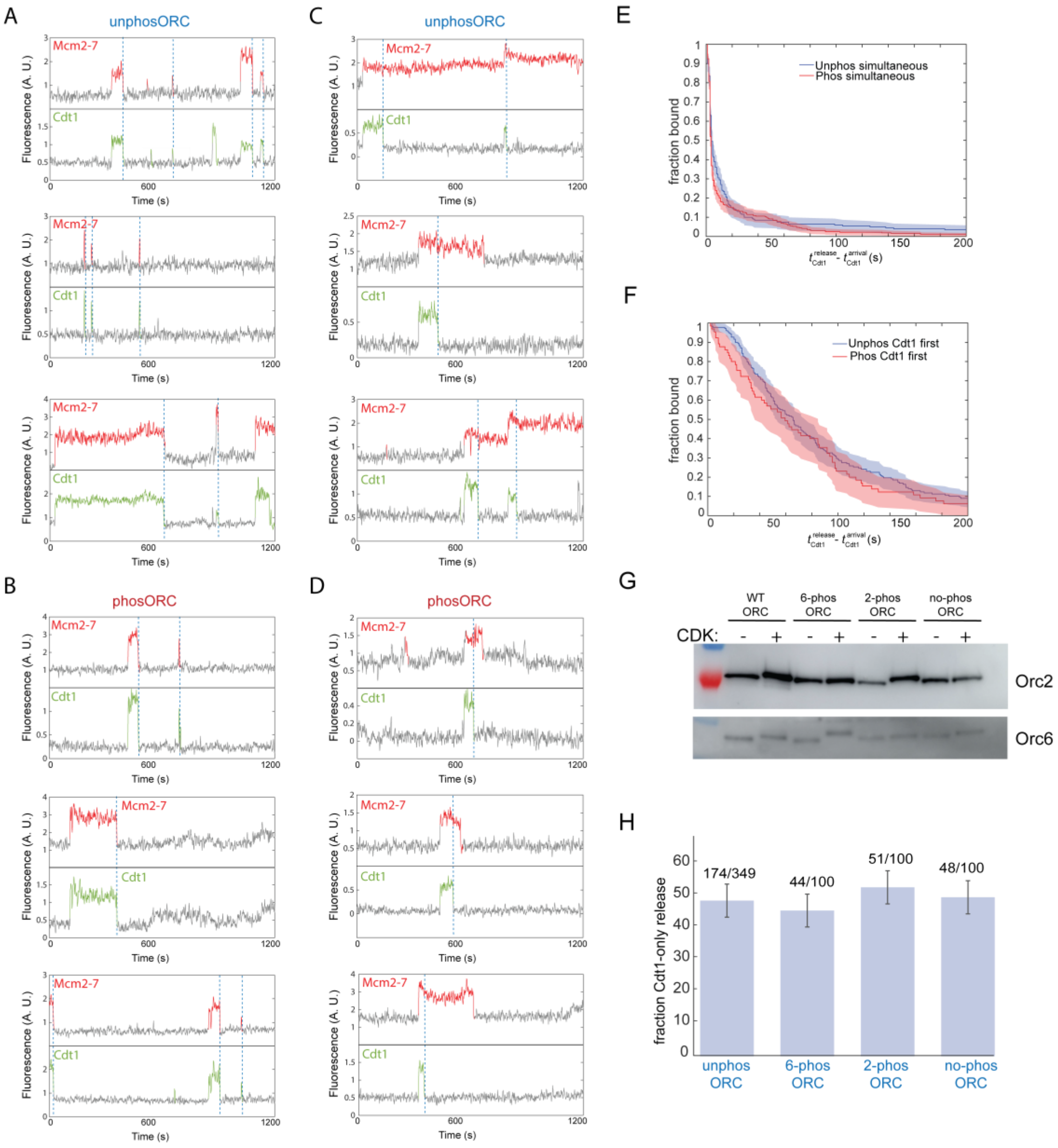


**Supplementary Figure 3. Additional Mcm2-7/Cdt1 fluorescence intensity traces, Cdt1 dwell-time distributions, and non-CDK-treated ORC phospho-site mutant controls.**

**A,B.** Additional example records showing non-productive Mcm2-7$^{4Sort\text{-}JF646}$ (red) and Cdt1$^{Sort\text{-}549}$ (green) simultaneous dissociation (dashed lines) from a single DNA (left pathway in Fig. 5A) for unphosORC (A) or phosORC (B). Plotted as in Fig. 5B.

**C,D.** Additional example records showing potentially productive Mcm2-7$^{4Sort\text{-}JF646}$ (red) and Cdt1$^{Sort\text{-}549}$ (green) in which Cdt1 dissociates from DNA before the corresponding first Mcm2-7 (dashed lines; right pathway in Fig. 5A) in the presence of unphosORC (C) or phosORC (D). Plotted as in Fig. 5C.

**E.** The fraction of Cdt1 molecules that remain on DNA for events where Mcm2-7 and Cdt1 leave simultaneously in the presence of unphosORC (blue), or phosORC, (red) measured from the time of Cdt1 arrival. Shaded areas represent 95% confidence intervals.

**F.** The fraction of Cdt1 molecules that remain on DNA for events where Cdt1 leaves before the associated Mcm2-7 in the presence of unphosORC (blue), or phosORC, (red) measured from the time of Cdt1 arrival. Shaded areas represent 95% confidence intervals.

**G.** Western blot of Orc2 and Orc6 showing CDK-dependent band-size increases for wt, 6-phos, 2-phos, and no-phos ORC. WT-ORC shows a CDK-dependent shift in both Orc2 and Orc6, whereas the 6-phos, 2-phos, and no-phos mutants show only shifts in Orc6, Orc2, or no band shifts, respectively.

**H.** The percentage (± SEM) of first Cdt1 dissociation events that followed the productive pathway (Cdt1 release without Mcm2-7 release) by unphosORC and the unphosphorylated (non-CDK-treated) ORC phospho-site mutants.

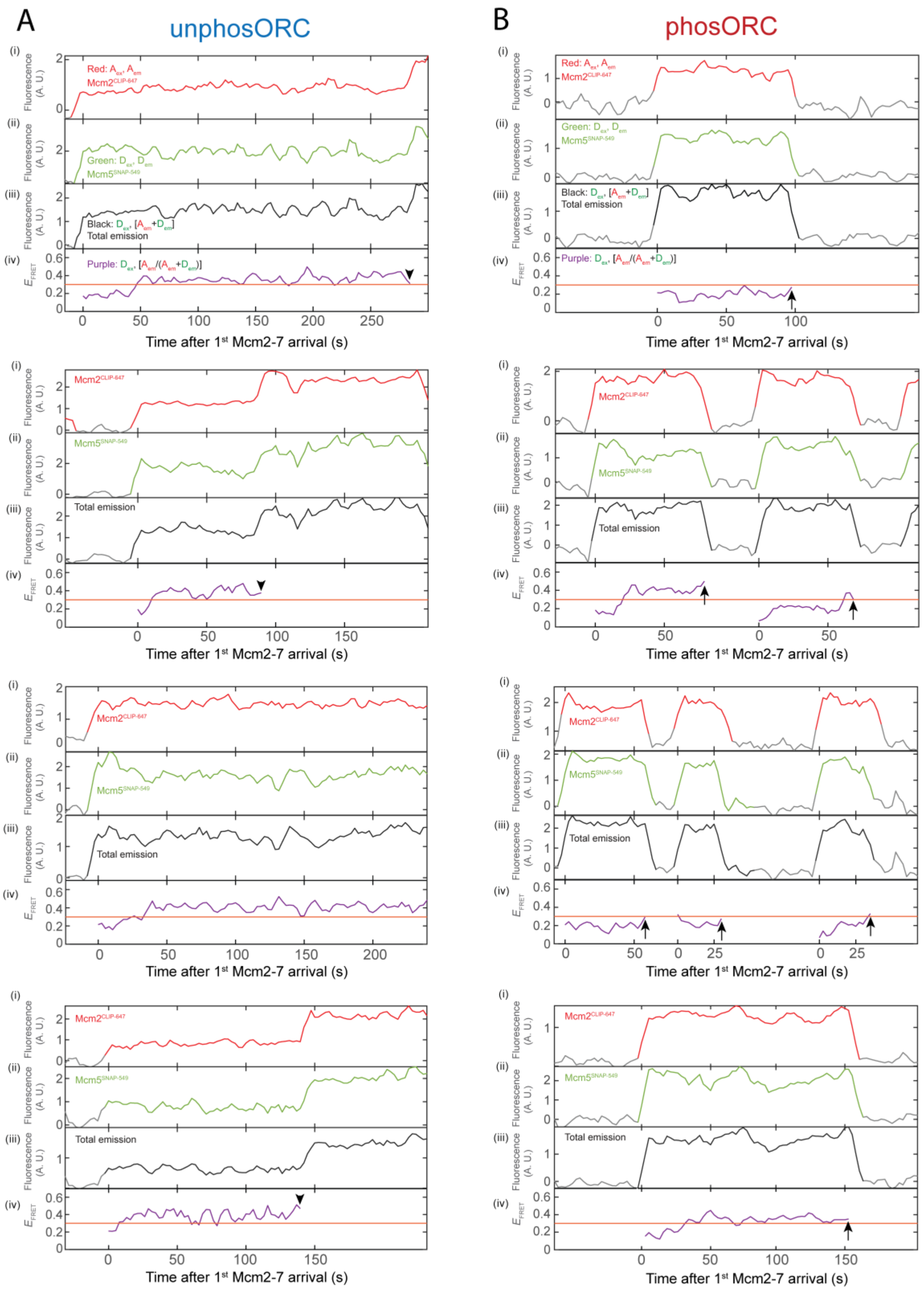


**Supplementary Figure 4. Additional Mcm2-7$^{25FRET}$ ring-closing FRET records from individual DNA molecules.**

**A.** Additional example records of Mcm2-7$^{25FRET}$ association with a single DNA molecule in the presence of unphosORC. Panels are as described in Fig. 6B. $E_{FRET}$ is shown only during intervals when fluorescence from both labeled subunits was present, or until a 2$^{nd}$ Mcm2-7 arrived (marked with arrowhead).

**B.** Additional example records of Mcm2-7$^{25FRET}$ association with a single DNA molecule in the presence of phosORC. Panels are as described in Fig. 6B. 1$^{st}$ Mcm2-7 release events are marked with arrow.

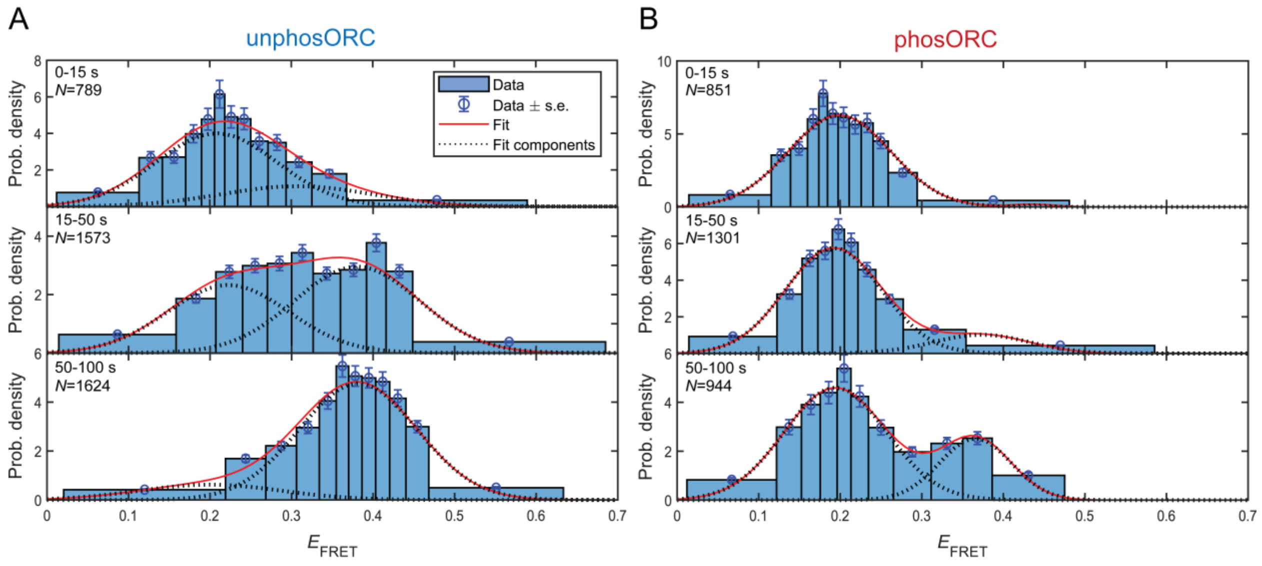


C

| Time interval (s) | Parameter | Value ± S.E. |
|---|---|---|
| 0-15 | $P_{low}$ | 0.7 ± 0.6 |
| | $\mu_{low}$ | 0.21 ± 0.02 |
| | $\sigma_{low}$ | 0.07 ± 0.01 |
| | $\mu_{high}$ | 0.31 ± 0.02 |
| | $\sigma_{high}$ | 0.09 ± 0.05 |
| 15-50 | $P_{low}$ | 0.4 ± 0.1 |
| | $\mu_{low}$ | 0.22 ± 0.02 |
| | $\sigma_{low}$ | 0.07 ± 0.01 |
| | $\mu_{high}$ | 0.38 ± 0.02 |
| | $\sigma_{high}$ | 0.08 ± 0.01 |
| 50-100 | $P_{low}$ | 0.14 ± 0.09 |
| | $\mu_{low}$ | 0.20 ± 0.06 |
| | $\sigma_{low}$ | 0.09 ± 0.02 |
| | $\mu_{high}$ | 0.38 ± 0.01 |
| | $\sigma_{high}$ | 0.07 ± 0.01 |

D

| Time interval (s) | Parameter | Value ± S.E. |
|---|---|---|
| 0-15 | $P_{low}$ | 0.99 ± 0.01 |
| | $\mu_{low}$ | 0.20 ± 0.004 |
| | $\sigma_{low}$ | 0.06 ± 0.003 |
| | $\mu_{high}$ | 0.43 ± 0.02 |
| | $\sigma_{high}$ | 0.03 ± 0.02 |
| 15-50 | $P_{low}$ | 0.85 ± 0.03 |
| | $\mu_{low}$ | 0.19 ± 0.004 |
| | $\sigma_{low}$ | 0.06 ± 0.003 |
| | $\mu_{high}$ | 0.37 ± 0.02 |
| | $\sigma_{high}$ | 0.06 ± 0.01 |
| 50-100 | $P_{low}$ | 0.72 ± 0.04 |
| | $\mu_{low}$ | 0.19 ± 0.01 |
| | $\sigma_{low}$ | 0.06 ± 0.01 |
| | $\mu_{high}$ | 0.37 ± 0.01 |
| | $\sigma_{high}$ | 0.04 ± 0.01 |

**Supplementary Figure 5. Distributions of Mcm2-7$^{25FRET}$ ring-closing $E_{FRET}$ values.**

**A.** Probability-density histograms of unphosORC-directed Mcm2-7$^{25FRET}$ $E_{\mathrm{FRET}}$ values (excluding outliers < 0.1 or > 0.7, 0.7% of data) for the indicated time intervals after 1$^{st}$ Mcm2-7 arrival. Each histogram was fit with two-component Gaussian mixture models (lines). Note: *N* values represent number of data points within each time-slice, not the number of Mcm2-7s (this number is given in Fig. 6C).

**B.** Probability-density histograms of phosORC-directed Mcm2-7$^{25FRET}$ $E_{\mathrm{FRET}}$ values (excluding outliers < 0.1 or > 0.7, 1.8% of data) for the indicated time intervals after 1$^{st}$ Mcm2-7 arrival. Each histogram was fit with a two-component Gaussian mixture models (lines).

**C.** Fit parameters for the unphosORC-directed Mcm2-7$^{25FRET}$ $E_{\mathrm{FRET}}$ values. Values are reported for each time interval and were determined separately for each distribution.

**D.** Fit parameters for phosORC-directed Mcm2-7$^{25FRET}$ $E_{\mathrm{FRET}}$ values. Values are reported for each time interval and were determined separately for each distribution.

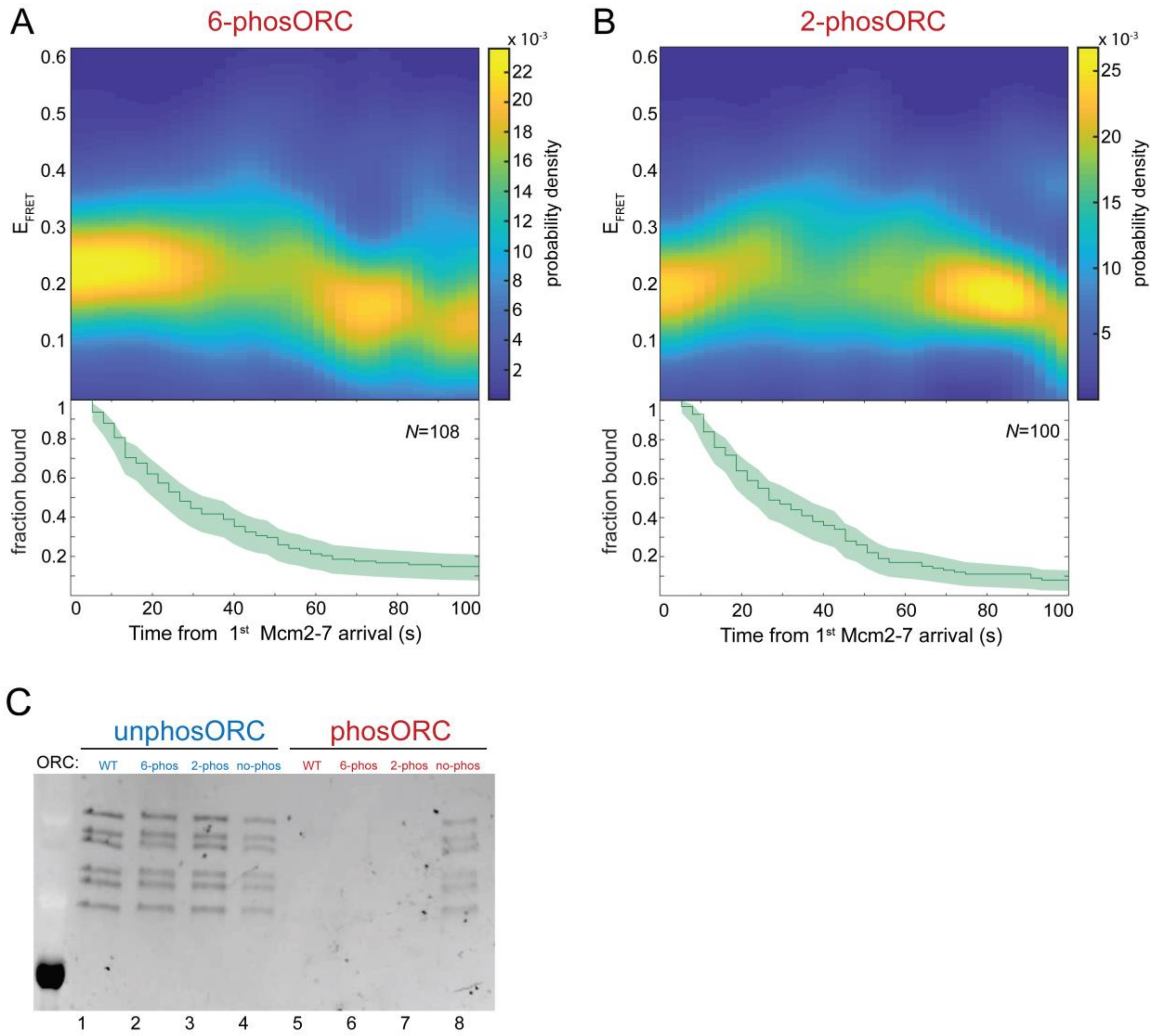


**Supplementary Figure 6. ORC with only the ORC6 subunit phosphorylated or only the ORC2 subunit phosphorylated inhibits Mcm2-7 ring closing and loading.**

**A.** Heat map of $E_{FRET}$ values vs. time after first Mcm2-7$^{25FRET}$ binding for $N$ = 108 DNA-bound 1$^{st}$ Mcm2-7 molecules in a reaction using 6-phosORC (top). Data were selected and plotted as described in Fig. 6C. The fraction of such complexes remaining at each time point is plotted (bottom).

**B.** Heat map of $E_{FRET}$ values vs. time after first Mcm2-7$^{25FRET}$ binding for $N$ = 100 DNA-bound 1$^{st}$ Mcm2-7 molecules in a reaction using 2-phosORC (top). The data were selected and plotted as described in Fig. 6C. The fraction of such complexes remaining at each time point is plotted (bottom).

**C.** Bulk helicase-loading assays comparing the loading of WT ORC (lanes 1 and 5), 2-phosORC (lanes 2 and 6), 6-phosORC (lanes 3 and 7) and no-phosORC (lanes 4 and 8) with and without phosphorylation. A high-salt wash (500 mM NaCl) was used to remove all helicase-loading intermediates from DNA.

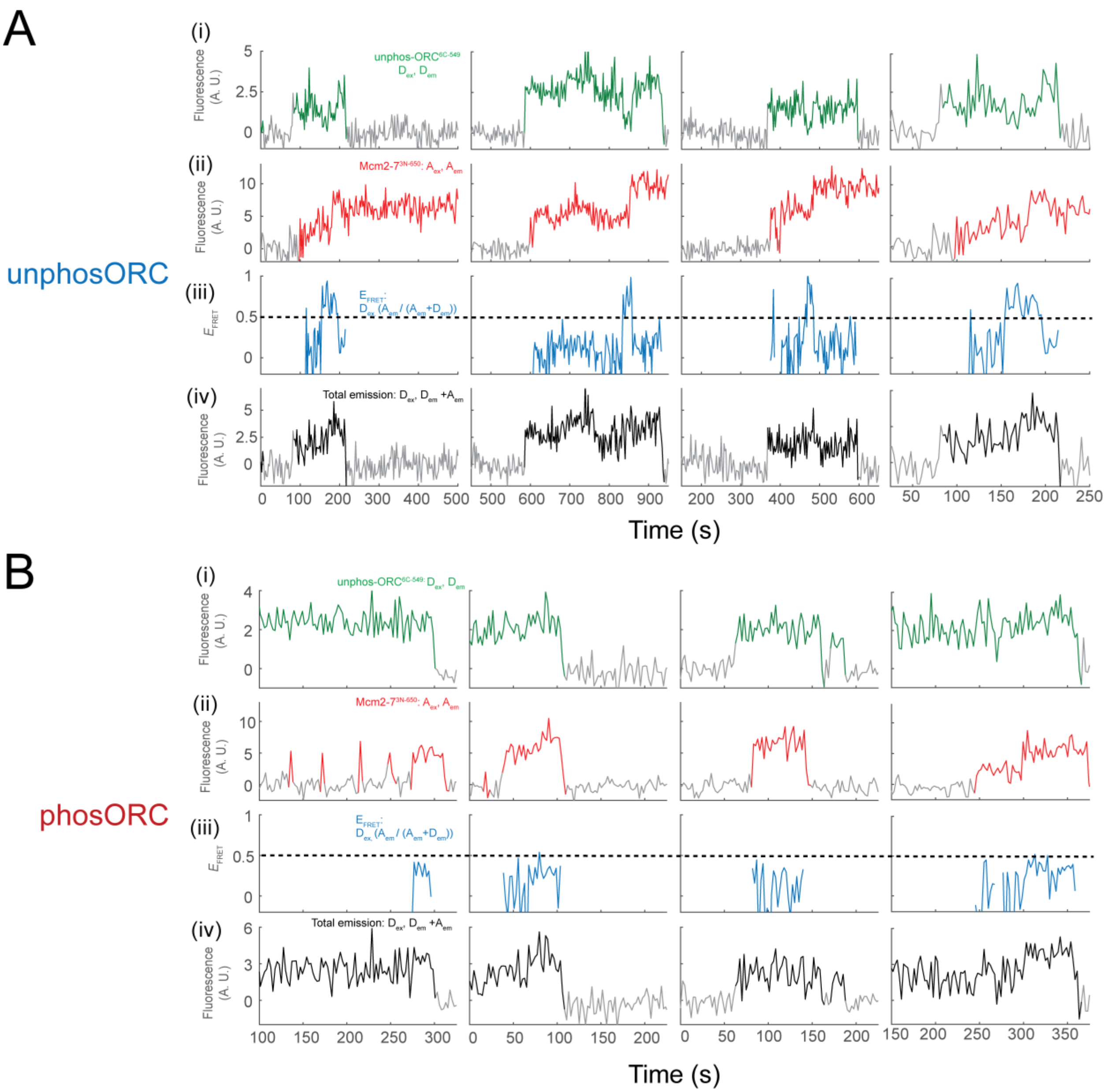


**Supplementary Figure 7. Additional records from individual DNA molecules for single-molecule MO-complex formation FRET assay.**

**A.** Additional example records of MO formation assays monitoring unphosORC$^{6C\text{-}549}$ and Mcm2-7$^{3N\text{-}650}$ association with a single DNA molecule. Panels are as described in Fig. 7B.

**B.** Additional example records of MO formation assays monitoring phosORC$^{6C\text{-}549}$ and Mcm2-7$^{3N\text{-}650}$ association with a single DNA molecule. Panels are as described in Fig. 7B.

**Supplementary Table 1: Fit parameters for Mcm2-7 binding to ORC-Cdc6-DNA.**

| ORC state | $A_f$ | $k_a$ ($\times 10^{-3}$ $s^{-1}$) | $k_{n,s}$ ($\times 10^{-3}$ $s^{-1}$) |
|---|---|---|---|
| unphosORC | 0.82 ± 0.02 | 4.7 ± 0.3<br>(N=466 DNA) | 0.09 ± 0.01<br>(N=625 non-DNA locations) |
| phosORC | 0.68 ± 0.03 | 2.0 ± 0.2<br>(N=612 DNA) | 0.09 ± 0.01<br>(N=526 non-DNA locations) |

Fits are for a single exponential binding model adjusted to remove the contribution from non-specific background DNA binding. Kinetic model is described in the Methods.

$k_a$ is the apparent first-order rate constant for Mcm2-7/Cdt1 binding to DNA. $k_{n,s}$ is the apparent first order rate constant for Mcm2-7/Cdt1 binding to controlnon-DNA sites (ie. non-specific binding to the surface of the slide). $A_f$ represents the "active fraction" of DNA molecules capable of recruiting Mcm2-7/cdt1.

**Supplementary Table 2: Yeast strains used in this study.**

| **Strain name** | **Expresses** | **Strain details** | **Study** |
|---|---|---|---|
| ySDORC | Wild-type ORC | *MATa ade2-1 ura3-1 his3-11,15 trp1-1 leu2-3,112 can1-100 bar1::hyg pep4::kanMX TRP1::pJF18 (GAL1,10 ORC5-opt ORC6-opt) HIS3::pJF17 (GAL1,10 ORC3-opt ORC4-opt) URA3::pJF19 (GAL1,10 CBP-TEV ORC1-opt ORC2-opt)* | Frigola et al., 2013 |
| yAA01 | 6-phosORC | *MATa ade2-1 ura3-1 his3-11,15 trp1-1 leu2-3,112 can1-100 bar1::hyg pep4::kanMX TRP::pJF18 (GAL1,10 ORC5-opt ORC6-opt) HIS3::pJF17 (GAL1,10 ORC3 ORC4) URA3::pAA04 (GAL1,10 Ubi-GGG-3x-Flag-ORC1 +ORC2-6Ala) ORC2-V5 (NatMX)* | This study |
| yAA02 | 2-phosORC | *MATa ade2-1 ura3-1 his3-11,15 trp1-1 leu2-3,112 can1-100 bar1::hyg pep4::kanMX TRP1::pAA02 (GAL1,10 ORC5-opt ORC6-opt-4Ala) HIS3::pJF17 (GAL1,10 ORC3-opt ORC4-opt) URA3::pAA04 (GAL1,10 Ubi-GGG-3x-Flag-ORC1-opt ORC2-opt) ORC6-V5 (HphMX)* | This study |

| | | | |
|---|---|---|---|
| yAA03 | no-phosORC | *MATa ade2-1 ura3-1 his3-11,15 trp1-1 leu2-3,112 can1-100 bar1::hyg pep4::kanMX TRP1::pAA02 (GAL1,10 ORC5-opt ORC6-opt-4Ala) HIS3::pJF17 (GAL1,10 ORC3-opt ORC4-opt) URA3::pAA04 (GAL1,10 Ubi-GGG-3x-Flag-ORC1-opt ORC2-opt-6Ala ORC6-V5 (HphMX)* | This study |
| yST166 | MCM2-7$^{4SNAP}$ and Cdt1$^{Sort}$ | *ade2-1 trp1-1 leu2-3,112 his3-11,15 ura3-1 can1-100 bar1::HisG lys2::HisG pep4::unmarked HIS3::pSKM004 (GAL1,10-MCM2, Flag-MCM3) URA3::pALS3 (GAL1,10 UbSORT-Cdt1, GAL4) LYS2::pST022 (GAL1,10 SNAP-MCM4, MCM5) TRP1::pSKM003 (GAL1,10 MCM6, MCM7)* | Ticau et al., 2015 |
| yST180 | Mcm2-7$^{4Sort}$ and Cdt1 | *ade2-1 trp1-1 leu2-3,112 his3-11,15 ura3-1 can1-100 bar1::HisG lys2::HisG pep4::unmarked TRP1::pSKM003 (GAL1,10-MCM6,MCM7) URA3::pALS1(GAL1,10-Cdt1,GAL4) HIS3::pSKM004(GAL1,10-MCM2,Flag-MCM3) LYS2::pST034 (GAL1,10 UbiSORT-MCM4,MCM5)* | Gupta et al., 2021 |
| yST229 | Mcm2-7$^{25FRET}$ | *ade2-1 trp1-1 leu2-3,112 his3-11,15 ura3-1 can1-100 bar1::HisG lys2::HisG pep4::unmarked TRP1::pSKM003 (GAL1,10-MCM6,MCM7) URA3::pALS1(GAL1,10-Cdt1,GAL4) HIS3::pST058 (GAL1,10 MCM2-721-CLIP,Flag-MCM3) LYS2::pST057 (GAL1,10 MCM4, MCM5-591-SNAP)* | Ticau et al., 2017 |
| yST103 | Cdt1$^{Sort}$ | *ade2-1 trp1-1 leu2-3,112 his3-11,15 ura3-1 can1-100 bar1::HisG lys2::HisG pep4::unmarked URA3::pST013 (GAL1,10 UbSORT-Cdt1-Flag)* | Gupta et al., 2021 |
| ySG39 | ORC$^{6C-549}$ | *ade2-1 trp1-1 leu2-3,112 his3-11,15 ura3-1 can1-100 bar1::HisG lys2::HisG pep4::unmarked LYS2::pSKM002 (GAL1,10 MCM4, MCM5) TRP1::pSKM003 (GAL1,10 MCM6, MCM7) HIS3::pSG13 (GAL1, 10 MCM2, Flag-TEV-GG-MCM3)* | Gupta et al., 2021 |
| ySG24 | Mcm2-7$^{3N-650}$ | *ade2-1 trp1-1 leu2-3,112 his3-11,15 ura3-1 can1-100 bar1::hisG lys2::HisG pep4::unmarked LYS2::pSKM002 (GAL1,10 MCM4, MCM5) TRP1::pSKM003 (GAL1,10 MCM6, MCM7) HIS3::pSG13 (GAL1,10 MCM2, Flag-TEV-GG-MCM3)* | Gupta et al., 2021 |

| | | | |
|---|---|---|---|
| ySG60 | 6-phos ORC$^{6C-549}$ | *MATa ade2-1 ura3-1 his3-11,15 trp1-1 leu2-3,112 can1-100 bar1::hyg pep4::kanMX HI3::pJF17 (GAL1,10 ORC3-opt, ORC4-opt) TRP1::pAZ63 (Gal1,10 ORC5-opt, ORC6-opt-C-LPETGG) URA3::pSG52 (Gal1,10 CBP-ORC1-opt, ORC2-opt-6A)* | This study |
| ySG61 | 2-phos ORC$^{6C-549}$ | *MATa ade2-1 ura3-1 his3-11,15 trp1-1 leu2-3,112 can1-100 bar1::hyg pep4::kanMX HIS3::pJF17 (GAL1,10 ORC3-opt, ORC4-opt) TRP1::pSG53 (GAL1,10 ORC5-opt, ORC6-opt-4A-C-LPETGG) URA3::pJF19(GAL1,10 CBP-ORC1-opt, ORC2-opt) ORC6-V5 (HphMX)* | This study |
| ySK119 | Clb5-Cdk1 | *ade2-1 trp1-1 leu2-3,112 his3-11,15 ura3-1 can1-100 bar1::HisG lys2::HisG pep4::unmarked URA3::GAL1,10 Δ2-95-CLB5-Flag CDC28-His* | Looke et al., 2017 |

**Supplementary Table 3: Plasmids used in this study**

| **Plasmid name** | **Construct encodes** | **Description** | **source** |
|---|---|---|---|
| pSKM033 | Flag-Cdc6 | pGEX- GST-PP-FLAG-Cdc6 | Kang et al., 2014 |
| pET-GSS-Cdc6 | Cdc6-sort | pET23b-GST-SUMO-GGG-Cdc6 | Ticau et al., 2015 |
| pAA02 | Orc5 and Orc6-4Ala | *GAL1,10 ORC5-opt*, *ORC6-opt-4Ala* | This study |
| pAA04 | UbSORT-FLAG-Orc1 and Orc2-6Ala | *GAL1,10 UBI-GGG-3xFlag-ORC1-opt, ORC2-opt-6Ala* | This study |
| pJF17 | Orc3 and Orc4 | *GAL1-,10 ORC3-opt, ORC4-opt* | Frigola et al., 2013 |
| pJF18 | Orc5 and Orc6 | *GAL1-,10 ORC5-opt, ORC6-opt* | Frigola et al., 2013 |
| pJF19 | CBP-Orc1 and Orc2 | *GAL1,10 ORC1-opt ORC2-opt* | Frigola et al., 2013 |
| pSG52 | CBP-Orc1 and Orc2-6Ala | *GAL1,10 CBP-ORC1-opt, ORC2-opt-6Ala* | This study |
| pSG53 | Orc5 and Orc6-4Ala-CLPETGG | *GAL1,10 ORC5-opt, ORC6-opt-4A-C-LPETGG* | This study |
| pAZ63 | | | |
| pSG13 | Mcm2 and Flag-TEV-GG-Mcm3 | *GAL1,10 MCM2, Flag-TEV-GG-MCM3* | Gupta et al., 2021 |
| pSKM002 | Mcm4 and Mcm5 | *GAL1,10 MCM4, MCM5* | Kang et al., 2014 |

| | | | |
|---|---|---|---|
| pSKM003 | Mcm6 and Mcm7 | *GAL1,10 MCM6, MCM7* | Kang et al., 2014 |
| pSKM004 | Mcm2 and Flag-Mcm3) | *GAL1,10 MCM2, Flag-MCM3* | Kang et al., 2014 |
| pST022 | SNAP-Mcm4 and Mcm5 | GAL1,10 SNAP-MCM4, MCM5 | Ticau et al., 2015 |
| pALS1 | Cdt1 and Gal4 | *GAL1,10 CDT1, GAL4* | Kang et al., 2014 |
| pALS3 | UbSORT-Cdt1 and Gal4 | *GAL1,10 UbSORT-CDT1, GAL4* | Ticau et al., 2015 |
| pST034 | UbiSORT-Mcm4 and Mcm5 | *pST034 GAL1,10 UbiSORT-MCM4, MCM5* | Ticau et al., 2015 |
| pST013 | UbSORT-Cdt1-Flag | *GAL1,10-UbSORT-CDT1-Flag* | Gupta et al., 2021 |
| pST058 | Mcm2-721-CLIP and Flag-Mcm3 | *GAL1,10 MCM2-721-CLIP, Flag-MCM3* | Ticau et al., 2017 |
| pST057 | Mcm4 and Mcm5-591-SNAP | *GAL1,10 MCM4, MCM5-591-SNAP* | Ticau et al., 2017 |
| pGEX-Sic1 | GST-Sic1 | *pGEX-4T-2-GST-Sic1* | Heller et al. 2011 |
